# A little feat in translation: miniaturized tRNAs in nematode and arachnid mitochondria

**DOI:** 10.64898/2026.06.12.731814

**Authors:** Josefine Gnauck, Iuliia Ozerova, Tim Kolberg, Sarah von Löhneysen, Heike Betat, Philipp H. Schiffer, Ina Schaefer, Peter Stadler, Mario Mörl

## Abstract

Transfer RNAs carry a highly conserved, cloverleaf-like secondary structure, which is essential to fulfill their task as adaptor molecules in translation. However, in mitochondria of metazoans, tRNA molecules were identified that deviate from this consensus structure and lack either the D- or T-arm, or, in some extreme cases, even both arms. These deviations are predominantly found in nematodes and arachnids, where in many cases the entire set of 22 mt tRNA genes are predicted to encode for such aberrant tRNAs or where even the complete loss of tRNA genes is proposed. Due to this unusual composition, we analyzed and characterized the mt tRNA pool of one representative of both groups. We identified the whole set of mt tRNAs and reannotated several tRNAs that differ significantly from previous genome-based predictions. In some cases, the sequence reads indicate putative tRNA editing events, showing that predictions exclusively based on mitogenome data have only a limited reliability. Our data also provide first insights into the modification pattern of such hairpin-like tRNAs.

## Introduction

Non-coding RNAs are essential modulators of a wide range of functions in the cell. To this end, not only sequence motifs, but also secondary and tertiary structure are crucial for RNA functionality. A prominent example for the interplay between structure and functionality are transfer RNAs (tRNAs). These molecules exhibit a cloverleaf-like secondary and an L-shaped tertiary structure to fulfill their task as adapters in translation. Accordingly, the tRNA structure is highly conserved in all domains of life. It consists of an acceptor stem that carries the indispensable 3’-CCA triplet serving as the amino acid attachment site, an anticodon arm that presents the anticodon for binding to the corresponding mRNA codon, a D- and a T-arm involved in tertiary structure formation and interaction with translation factors and maturation enzymes ^1^. In addition to cytosolic tRNAs encoded in the nuclear genome, mitochondria and chloroplasts carry their own tRNA set encoded in their genome. The metazoan mitochondrial tRNA pool consists of 22 tRNAs, most of them adopting the typical canonical structure ^2^. In human and bovine mitochondria, however, it was shown that tRNA^Ser^(AGY) lacks the D-arm, representing the first deviation from the conserved shape ^3,4^. Soon, more tRNAs with aberrant structures were discovered in mitochondria of metazoans, and these tRNAs lack either the D-, the T-arm or even both arms that are replaced by single-stranded connector elements ^5–10^. A systematic analysis of all available metazoan mitogenomes predicts that about 50% of the genomes encode for at least one deviating tRNA structure ^11^. Especially in the groups of arachnids and nematodes truncated tRNAs seem to accumulate, and in some of these organisms, all 22 mt tRNAs display an aberrant structure. These findings demonstrate that truncated mt tRNAs do not represent an exotic or exceptional feature but rather are common in metazoans. Nonetheless, these results rely exclusively on computational predictions. Even though specific annotation algorithms were developed to take the reduced tRNA size into account ^12,13^, the prediction of armless, hairpin-like tRNAs solely based on mitogenome data remains challenging. While for Enoplea nematodes a specific algorithm could predict the whole set of highly truncated tRNAs with only slight deviations ^6^, such a tool is not available for other phyla. Within the last two decades, the availability of mitogenomes of the superorder of Acari (arachnids) increased dramatically and resulted in further computer-based predictions of truncated tRNAs, although with contradicting results ^5,14–18^. In species like *Steganacarus magnus*, *Sarcoptes scabiei*, *Tyrophagus longior* and *Tyrophagus putrescentiae*, it was not possible to identify the whole set of 22 mt tRNAs and, consequently, a loss of these mt tRNA genes was proposed ^5,19,20^. A complicated case is demonstrated in *Steganacarus magnus*, where lost mt tRNAs genes were retrieved and existing mt tRNA genes were re-annotated ^5,14–18^. Hence, the question arises whether the predicted truncated tRNAs are actually transcribed and functional.

A first proof was given by Wende *et al*. ^10^, where three of nine predicted armless mt tRNAs of *Romanomermis culicivorax* (Enoplea) were verified at the RNA level. Further - coincidental - evidence emerged from the sequence analysis of mitochondrial tRNAs of the angiosperm *Silene vulgaris* that additionally contained the mitochondrial tRNA pool of the contaminating plant pest *Tetranychus urticae* (Acari), including several armless tRNAs ^21^. While these data represent clear evidence for the existence of truncated, armless tRNAs, further sequence analyses would (1) help improve prediction algorithms, (2) extend our understanding of functionality and structure of armless tRNAs and (3) allow the investigation of post-transcriptional editing and base modifications.

Here, we analyzed the mitochondrial tRNA pool of one representative species of the two phyla known to encode highly truncated mt tRNAs: the oribatid mite species *Steganacarus magnus* (*Sma*; arachnid) and the mermithid nematode species *Romanomermis culicivorax* (*Rcu*; Enoplea). As the sequencing of highly structured and modified tRNAs is challenging, we used a combination of different library preparation methods. In LOTTE-Seq, we used a specific tRNA ligation reaction, where a hairpin adapter complementary to a tRNAs 3’-NCCA end is fused to its target ^22^. This method was combined with a demethylation step to reduce RT stops ^23^, as well as an efficient circularization of the cDNA instead of a second adapter ligation ^24^. We identified the entire mt tRNA pool for both organisms, and in *R. culicivorax*, the identified transcripts correctly match the predictions ^6^. In *S. magnus*, however, the mt tRNA sequences revealed strong deviations from most predictions ^5,14–18^. Furthermore, expression of overlapping sense and antisense tRNA genes was identified, as described for *T. urticae* ^21^. The analysis also revealed 3‘-editing events for several tRNAs as well as the first modification patterns for armless tRNAs. So far structural data of armless tRNAs only exist for *in vitro* substrates ^25^. Hence, we investigated the secondary structure of native armless tRNAs by Led-Seq ^24^, showing that these tRNAs have a hairpin-like structure highly similar to that of the corresponding *in vitro* transcripts.

## Results and Discussion

### A combinatorial approach for efficient tRNA sequencing

High-throughput sequencing of tRNAs remains a challenge, as their stable structure and high frequency of modifications interfere with reverse transcription. Moreover, mitochondrial tRNAs only present a small proportion of the total tRNA pool of a cell and are therefore underrepresented in sequencing libraries^26^. In the last decade, several methods for sequencing library construction have been developed that focus on overcoming obstacles of reverse transcription ^22,23,26–28^. The introduction of the demethylating enzyme AlkB in ARM-Seq and DM-TGIRT-Seq made it possible to remove modifications m^1^A, m^3^C and m^1^G, eliminating these road blocks of reverse transcription ^23,28^. To tackle the structural barrier for cDNA synthesis, the thermostable group II intron reverse transcriptase (TGIRT) was implemented ^28^, which also allows the quantification of modifications by an adjusted protocol ^26^. The template-switching activity of TGIRT allows to bypass the adapter ligation step but requires purification of tRNAs from total RNA to produce only tRNA-derived reads. In YAMAT-Seq and LOTTE-Seq, a tRNA specific ligation reaction was introduced that allows the isolation of mature tRNAs by their 3’-CCA-end out of a total RNA preparation ^22,27^.

Nucleic acid material obtained from small, non-model/laboratory organisms is a limiting factor for tRNA analyses. Therefore, we combined the advantages of LOTTE-Seq and ARM-Seq and introduced an alternative cDNA circularization step (Fig. 1) ^24^. This procedure allowed us to produce a sequencing library out of a very limited amount of total RNA without prior selection of tRNAs. Furthermore, the comparison of AlkB-treated and untreated aliquots was used as a read-out to identify base methylation patterns, as these modifications cause RT-stops in the untreated samples. For unambiguous genome mapping and tRNA identification, only sequencing reads longer than 8 nt with CCA end, UMIs (unique molecular identifier) and Illumina-specific sequences were considered for further analysis. Depending on the individual library, this fraction consisted of 60% – 73% of total reads (Suppl. Fig. 1). A mapping summary of the mt tRNA sequences is depicted in Figure 2A and 2C.

**Figure 1:**
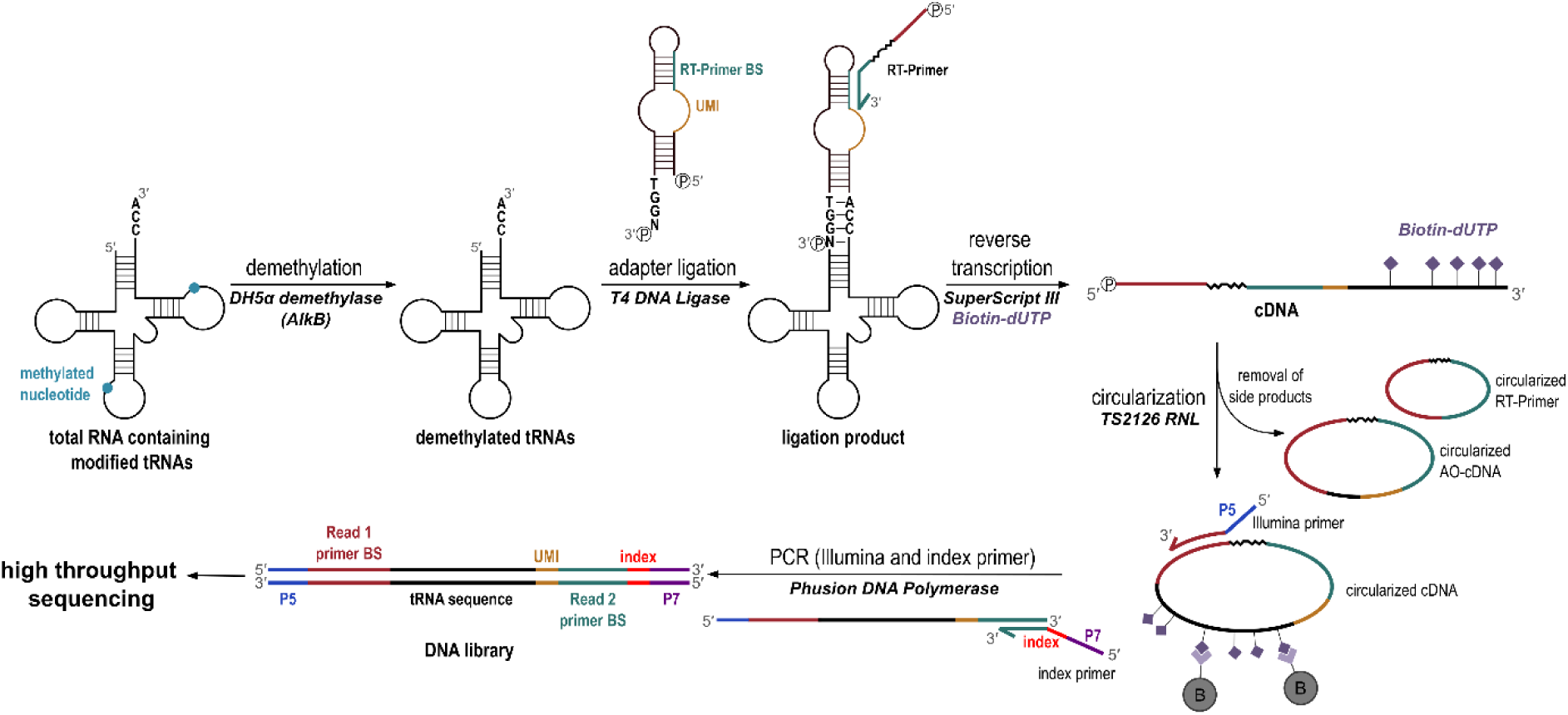
Combinatorial approach for tRNA sequencing - LOTTE-Seq and ARM-Seq. Methylations at nucleotides (green) are removed by AlkB demethylase treatment ^23^. In a second step tRNAs are specifically ligated to a hairpin adapter hybridizing to the CCA-end and the upstream located single-stranded discriminator position ^22^. To identify possible amplification artifacts, the hairpin adapter carries an 8 nt UMI (unique molecular identifier) sequence consisting of randomized T, C, and G positions (orange). Reverse transcription starts with an RT primer binding to the hairpin adapter and is carried out with biotinylated nucleotides (purple diamonds) for specific enrichment of reaction products. The obtained cDNA is circularized, and the streptavidin bead clean-up procedure removes non-specific byproducts like circularized RT primer without cDNA and circularization products containing adapter sequence only (AO-cDNA). The immobilized cDNA is used directly for subsequent PCR, where Illumina-compatible primers introduce the required Illumina sequences as well as library-specific indices ^24^.

**Figure 2:**
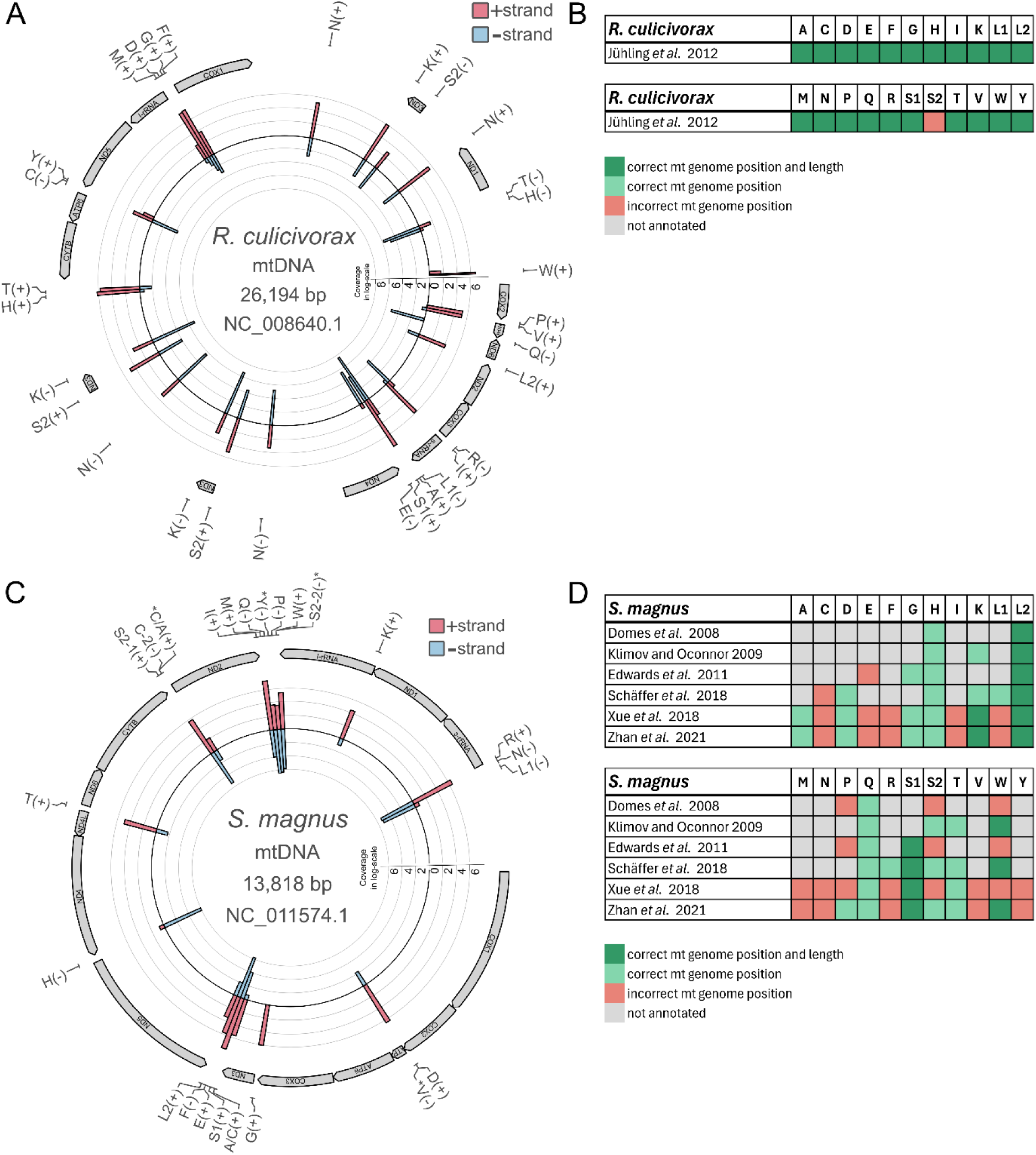
Mapping summary of tRNA sequences to the mitogenomes of *R. culicivorax* and *S. magnus.* Outer circle shows mt tRNA annotation retrieved from experimental data for *R. culicivorax* (**A**) and *S. magnus* (**C**). (+) and (-) symbols indicate sense or antisense orientation of the corresponding genes. tRNA identity is shown by the amino acid one-letter-code. Bars represent read numbers for (+)- (red) and (-) strand-encoded (blue) tRNAs. Low coverage tRNAs are indicated by an asterisk. (**B; D**) Bar diagrams for mt tRNA gene locations compared to computer-based predictions. (**B**) For *R. culicivorax*, only a single gene for tRNA^Ser2^ was not correctly predicted (red), while the rest of the tRNAs were correctly proposed in terms of position and transcript length (green), demonstrating the high reliability of the algorithm used by Jühling et al. ^6^. (**D**) In contrast, for *S. magnus*, size and genome position for only very few tRNA genes were correctly predicted in a series of publications (green), indicating the importance of characterizing these tRNAs at the transcription level ^5,14–18^.

### The set of mt tRNAs from *R. culicivorax* matches with prediction

In *R. culicivorax*, most of the identified mt tRNAs correlate with computational predictions with two D-armless, eleven T-armless and nine armless tRNAs ^6^ (Fig. 3). Only minor alterations were observed, where an additional (tRNA^Ala^, tRNA^Met^) or missing nucleotide at each end (tRNA^Ile^, tRNA^Phe^, tRNA^Cys^, tRNA^His^, tRNA^Lys^) was found. Compared to the prediction, tRNA^Ser2^ lacks two A residues within the T-arm (Fig. 3). As a result, this sequence aligns to a different mitogenome position (Fig. 2B). This suggests that, while the class-specific prediction algorithm for Enoplea yields excellent results, an analysis of transcriptomic data is necessary for the validation of mature tRNAs. Even though the additional or missing nucleotides at the 5’- and 3’-ends represent only minor changes, this should not be underestimated, as they affect the identity of the discriminator base and are therefore essential for the functionality of tRNAs ^29^.

**Figure 3:**
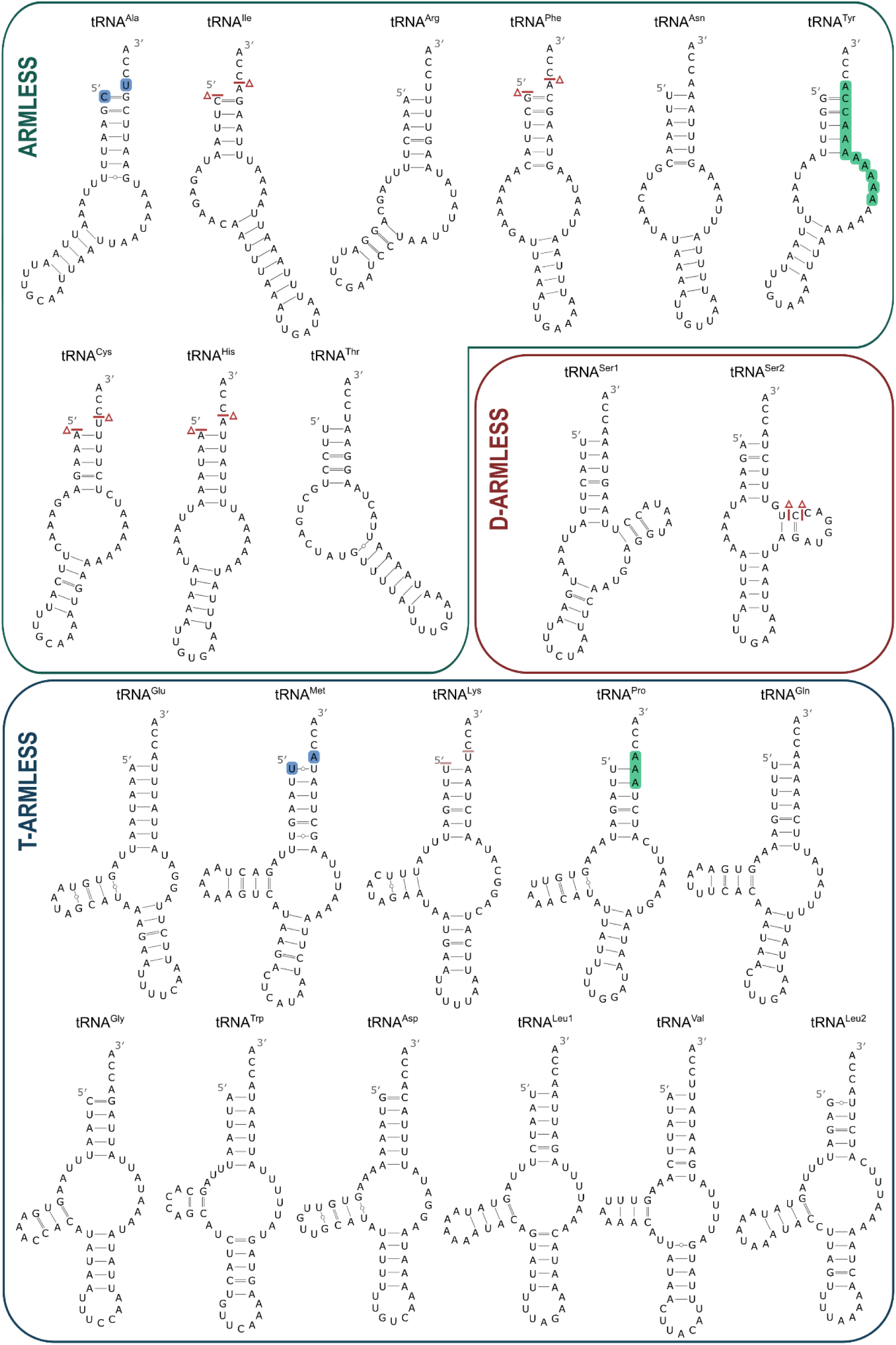
Secondary structure prediction of mt tRNAs from *R. culicivorax.* Secondary structures of the tRNAs identified by LOTTE-Seq were predicted by covariance models ^6^. Deviations from the sequences predicted by Jühling et al. are highlighted as follows: one additional nucleotide at each tRNA end (blue); missing nucleotide at each tRNA end or within the sequence (red delta symbol) and exchanged nucleotides due to editing (green).

Several non-Watson-Crick base pairs were identified in T-arm, acceptor arm as well as anticodon arm, including ten G-U pairs (tRNA^Ala^, tRNA^Thr^, tRNA^Glu^, tRNA^Lys^, tRNA^Pro^, tRNA^Asp^, tRNA^Val^, tRNA^Leu2^) and one U-U pair (tRNA^Met^). In particular, G-U pairs are a frequently observed tRNA feature and can serve as identity elements for aminoacyl-tRNA synthetases. In contrast, U-U pairs have not yet been identified in tRNAs. Due to stabilizing neighboring Watson-Crick base pairs, such U-U interactions can be part of a stable stem structure ^30–32^. Since the U1-U54 interaction represents the first base pair in the acceptor stem of tRNA^Met^, a stabilizing effect of the following U2-A53 pair may be negligible. For the initiator tRNA^Met^ of *E. coli*, it was shown that a less stable C1-A72 base pair is necessary for translation initiation ^33^. As this mismatch is a unique feature of prokaryotic initiator tRNAs, the mismatch U1-U54 in *R. culicivorax* mt tRNA^Met^ could be a remnant of endosymbiotic origin, as mitochondrial translation is also initiated by fMet-tRNA^Met 34^. The U1-U54 base pair instead of the C1-A72 found in prokaryotes could be a specific feature allowing a single mt tRNA^Met^ acting both as initiator as well as elongator tRNA. For mt tRNA^His^, the highly conserved G-1 position, an identity element for the corresponding aminoacyl-tRNA synthetase, was not observed ^35,36^. As it is not encoded in the mt genome of *R. culicivorax*, the obtained sequence probably represents a maturation intermediate. The fact that fully processed tRNA^His^ with G-1 is not found in the sequencing data can be explained by the tRNA-specific adapter hybridization that requires a single stranded CCA-end plus the adjacent discriminator base in the tRNA as a prerequisite for ligation to position 1 of the tRNA. Therefore, a G-1 poses a sterical problem, and adapter ligation is only possible with premature tRNA^His^ lacking G-1.

### Structure probing of native armless tRNAs shows hairpin-like structures

In a previous study, in-line probing experiments on *in vitro* transcribed mt tRNA^Ile^ and mt tRNA^Arg^ from *R. culicivorax* revealed a hairpin-like secondary structure ^25^. As these transcripts were accepted as substrates by the corresponding CCA-adding enzyme and elongation factor mt EF-Tu1 ^37,38^, they likely represent a functional form. To examine whether native mt tRNAs also fold into these structures, the mt tRNA preparation was subjected to Led-Seq analysis that is based on lead-induced cleavage at single-stranded positions and subsequent analysis by adapter ligation and deep sequencing ^24^.

Compared to the probing of canonical tRNAs, we observed a higher background signal for the examined mt tRNAs. This could be caused by the high AU content of the armless tRNAs, as A-U base pairs are more prone to fraying. This background signal, combined with the short length of the transcripts, leads to the appearance of more open structures than the actual RNAs might exhibit after implementing the probing data into the structure prediction pipeline of the Led-Seq approach. Therefore, we decided to utilize only the normalized probing signal (*S*) and its moving average to illustrate single-stranded regions. Hence, the probing signals only provide an approximation of the tRNA conformations in the data set.

As representative examples, Led-Seq probing of native mt tRNA^Ile^ and mt tRNA^Arg^ are shown in Fig. 4, exhibiting structures that correspond to those of their *in vitro* transcripts ^25^. The normalized probing signal (*S*) follows the RNAs secondary structure, where peaks are identified as positions being more susceptible to lead cleavage and therefore less likely to be involved in base-pairing. High signal intensities could be identified for the connector elements, the anticodon loop and for some positions in the acceptor stem, probably resulting from fraying events of these AU-rich regions. Applying a moving average through the single data points illustrates that the signal follows the proposed hairpin-like secondary structure (Fig. 4). Similar results were identified for other armless tRNAs of *R. culicivorax,* with some exceptions for mt tRNA^Cys^, mt tRNA^Phe^ and mt tRNA^Tyr^, where the anticodon stem shows an increased signal intensity (Supp. Fig. 2). Likewise, the D-armless mt tRNA^Ser1^ and mt tRNA^Ser2^ exhibit an ambiguous signal, which makes it difficult to assess their structure. T-armless tRNAs show a high signal intensity for the anticodon loop, the 3′-end and the respective connectors. The size-reduced D-arms also exhibit strong signals, which may be the result of the absence of tertiary interactions in the elbow region, leading to an increased flexibility in these arms (Supp. Fig. 2). Overall, despite some uncertainties, the normalized probe signal from native armless tRNAs supports the secondary structure models derived from *in vitro* transcripts and computer-based predictions. Nevertheless, it should be noted that *ex vivo* RNA structures may deviate from *in vivo* conformations to a certain extent, as salt concentrations, temperature, metal ions, interactions with proteins and cellular localization can have an impact on the structural organization ^39^. Hence, the shape of isolated tRNAs may not be as stable as within the cell.

**Figure 4:**
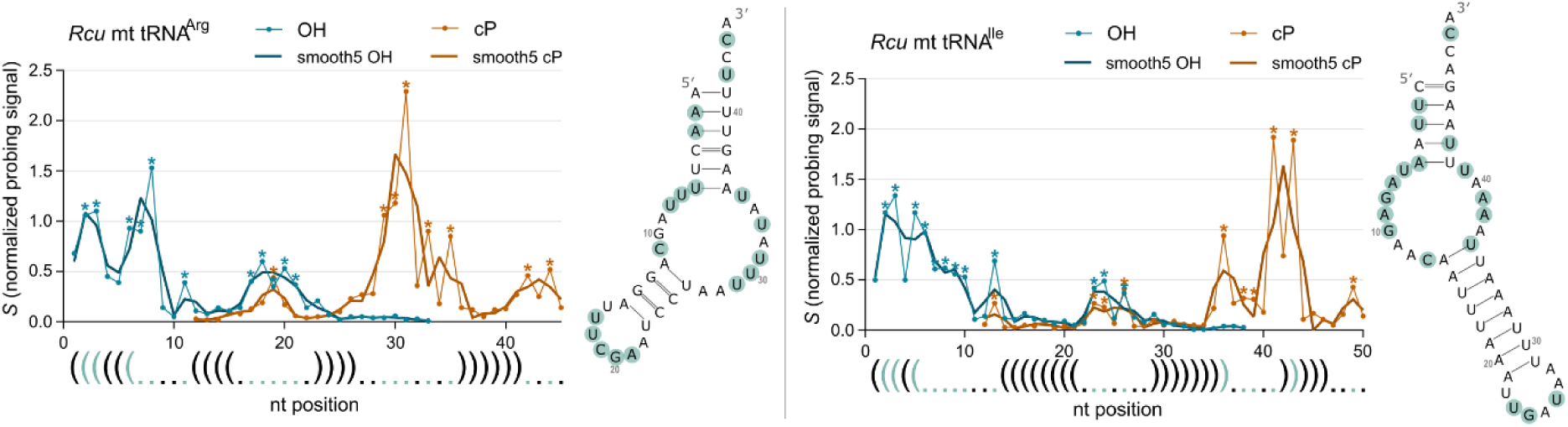
Led-Seq-derived structural data of native armless mt tRNAs of *R. culicivorax*. Diagrams show the normalized probing signal (S) at the different positions throughout the tRNA sequence. Signals from OH-libraries (representing 3’-fragments after lead-cleavage) are depicted in blue and signals from cP-libraries (5’-fragments) in orange. Signals that are more intense than their neighboring signals are marked with an asterisk and indicate a higher probability of the nucleotide being unpaired. Matching nucleotides are marked in the tRNA secondary structure with green circles. In addition, a moving average has been applied to the signal with a window size of 5 (smooth5 OH and smooth5 cP), which aligns with the depicted dot-bracket-notation of the structure. The probing signal of mt tRNA^Arg^ (left) and mt tRNA^Ile^ (right) follows the predicted hairpin-like structure. Background signals in the acceptor stem probably represent fraying events of the AU-rich stem.

### mt tRNAs from *S. magnus* are shorter than predicted

With an average length of 45 nucleotides, the mitochondrial tRNAs from *S. magnus* are shorter than anticipated. Secondary structure prediction of the obtained mt tRNAs was performed using RNAfold software of the Vienna RNA Package 2.0 ^40^ and resulted in 17 armless tRNAs, three T-armless tRNAs and two D-armless tRNAs, with the remaining D- and T-arms also exhibiting significant shortening (Fig. 5). A comparison of the transcriptome data with published genome positions indicates that about half of the tRNA genes were incorrectly annotated, and only four mt tRNAs were predicted with a length similar to that of the identified transcripts (Fig. 2D). This discrepancy is probably caused by the extremely reduced tRNA lengths and a high proportion of posttranscriptional editing events (see below). The difficulty of mt tRNA gene annotations solely based on *S. magnus* mitogenome data is demonstrated by the number of published re-annotations ^5,14–18^. In addition, the mt tRNAs show a high number of non-Watson-Crick base pairs (Fig. 5). In total, we observed 13 G-U base pairs (tRNA^Lys^, tRNA^Asn^, tRNA^Glu^, tRNA^Gln^, tRNA^His^, tRNA^Leu1^, tRNA^Ile^, tRNA^Pro^, tRNA^Ser2–2^), four U-U base pairs (tRNA^Asp^, tRNA^Gly^, tRNA^Arg^, tRNA ^Ala/Cys^) as well as one A-C base pair (tRNA^Met^), one G-A base pair (tRNA^Tyr^) and two A-A base pairs (tRNA^Val^, tRNA^Ala/Cys^). While G-U base pairs are frequently found in mt tRNAs and can serve as identity elements ^41^, the other non-canonical pairs are not common in tRNA stems. Yet, thermodynamic studies have shown that single mismatches can be found within a stable RNA duplex ^42^. For tRNA^Ser2^, we identified two candidates with identical anticodons and comparable read numbers (Fig. 5, grey box). Similarly, the identified sequences of tRNA^Ala^ and tRNA^Cys^ show ambiguous genome locations and structural foldings, respectively. At the annotated genome location for tRNA^Cys^, expression of both strands was detected. The (+)-strand transcript can adopt two alternative structures representing either tRNA^Cys^ or tRNA^Ala^ and has therefore been designated as tRNA^Cys/Ala^. The transcript mapping on the (-)-strand represents a tRNA^Cys^ candidate (designated as tRNA^Cys-2^). Furthermore, at the annotated genome location for tRNA^Ala^, expression of a transcript was detected that is also predicted to adopt two alternative structures, representing either a tRNA^Ala^ or a tRNA^Cys^ and is referred to as tRNA^Ala/Cys^.

**Figure 5:**
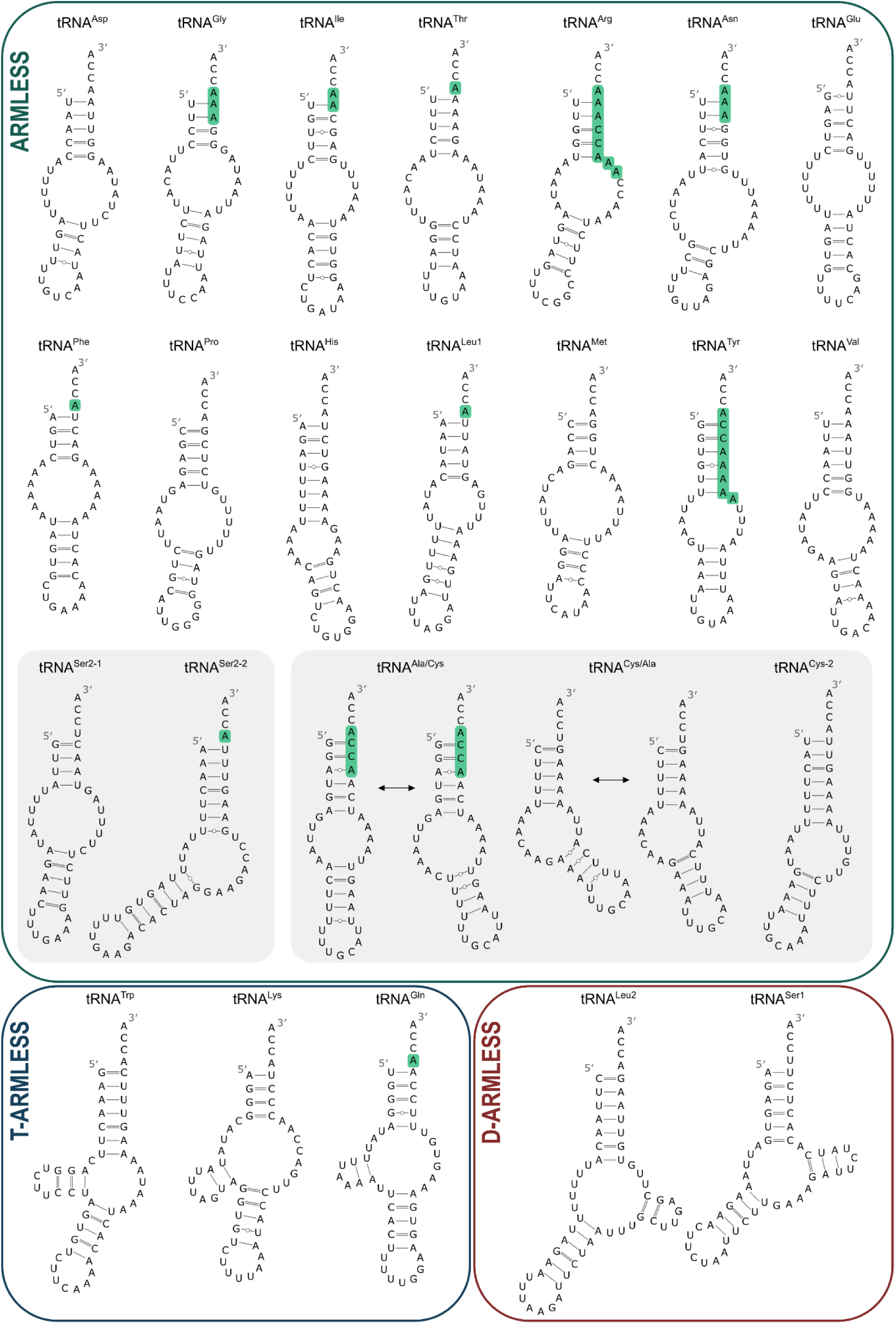
Secondary structure prediction of mt tRNAs from *S. magnus.* Secondary structures were predicted by RNAfold and manually adjusted to fit the typical tRNA structure with a 7 nt anticodon loop and an unpaired discriminator base at the 3’-end. tRNAs with more than one candidate structure or sequence are highlighted by a grey box. Edited nucleotides are marked in green.

As biological material of *S. magnus* is very limited, a Led-Seq approach of the tRNA preparation to clarify the structure of these transcripts was not feasible. Hence, the corresponding *in vitro* transcripts were investigated by in-line probing. The experiments confirmed that tRNA^Cys/Ala^ does only form a stable anticodon stem with a UGC anticodon (alanine) is presented, while the structure carrying a cysteine anticodon (GCA) contains rather weak A-A, A-C, and U-U interactions (Fig. 6A). It is therefore likely that this tRNA does not represent a tRNA for cysteine, but rather a tRNA for alanine. In contrast, tRNAs^Ala/Cys^ can form two stable anticodon stems presenting either anticodon UGC (alanine) or GCA (cysteine), and the in-line probing suggests that both conformations are possible (Fig. 6B). Both stems carry a non-canonical U-U base pair, but only the UGC anticodon stem contains a stabilizing C-G base pair. Hence, this transcript might also represent a tRNA^Ala^. When taking the number of sequence reads into account, the tRNA^Cys/Ala^ is highly underrepresented (102 reads), while tRNA^Ala/Cys^ is significantly more abundant (5,994). Further, the tRNA^Cys-2^ transcript shows a higher abundance, too (1,221 reads) (Suppl. Table 2B). According to the tRNA punctuation model of Ojala, metazoan mt transcription results in huge sense and antisense polycistronic precursors that are processed into individual mRNAs and rRNAs by the specific 5’- and 3’-cleavage events at tRNA or tRNA-like structures ^43^. Hence, the low abundant mt tRNA^Cys/Ala^ of *S. magnus* might represent such a tRNA-like processing signal of the (+)-strand rather than a functional tRNA. Since it resembles a tRNA structure, it was further processed into a CCA-carrying transcript that was caught by our CCA-hybridizing adapter oligonucleotide. Accordingly, tRNA^Ala/Cys^ is a bona-fide tRNA for alanine, while tRNA^Cys-2^ represents a true tRNA for cysteine.

**Figure 6:**
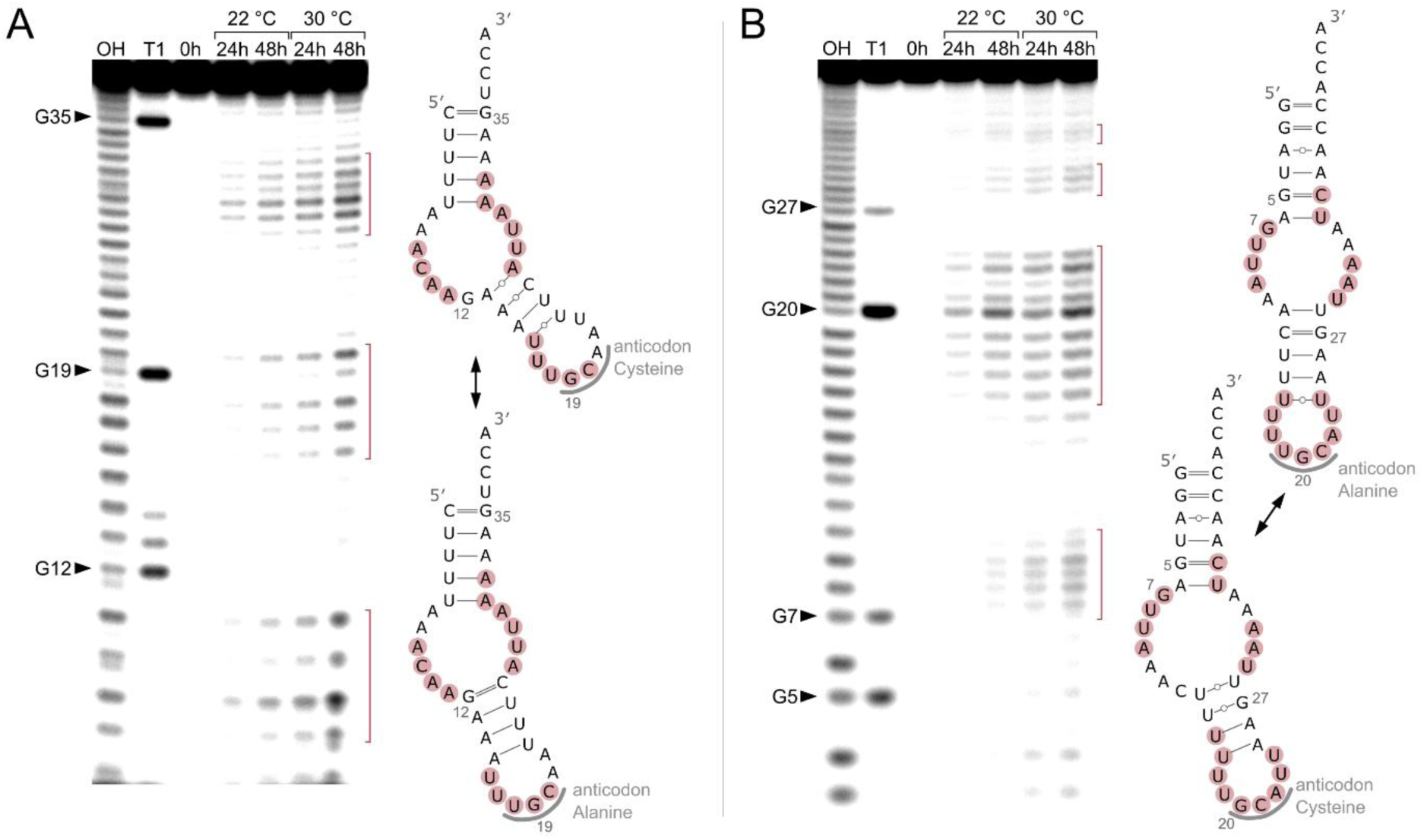
Structural data obtained through in-line probing of *in vitro* mt tRNAs from *S. magnus.* Probing signals were obtained at 22° C and 30° C. Identified single-stranded regions of mt tRNA^Ala/Cys^ (**A**) and tRNA^Cys/Ala^ (**B**) are depicted in the autoradiogram by red bars and mapped to the predicted secondary structures (red circles). Structure probing was performed in three independent experiments (n=3).

### Sense and antisense mt tRNA transcripts occur in both *S. magnus* and R. culicivorax

Further sense/antisense expression was detected for tRNA^Ile^, tRNA^Met^, tRNA^Gln^, tRNA^Arg^ and tRNA^Pro^ with a read number between 1% and 10 % of the actual transcript (Fig. 2C, Supp. Table 2B). Again, the antisense transcripts can form tRNA-like structures leading to CCA addition, so that the hairpin adapter can be ligated and the corresponding sequences appear in the LOTTE-Seq 2 analysis. These transcripts probably also serve as punctuation signals of the corresponding precursor transcript. In three cases, however, the transcription of (+)- and (-)-strands resulted in sense and antisense transcripts, both of which serve as functional tRNAs. This is the case for tRNA^Glu^/tRNA^Phe^, tRNA^Asp^/tRNA^Val^, and tRNA^Trp^/tRNA^Ser2–2^ (Supp. Table 2B). This feature is particularly striking as a single mutation within this region would affect both tRNAs, leading to an increased pathogenicity ^44^. Notably, mitochondrial dual-strand expression and punctuation has been previously described in *T. urticae* ^21^, indicating that this might be a common feature for the subclass of Acariformes and a remnant of the ancestral decoding system of life ^45^.

Similarly, several mt tRNA genes show antisense transcription in *R. culicivorax*, albeit with low expression levels (1-13%) compared to the sense transcripts representing tRNAs. Hence, it is very likely that these antisense transcripts also represent punctuation signals for primary transcript maturation.

### Mt tRNAs of both organisms are subject to 3**′**-editing

tRNA editing is a widespread feature in metazoan mitochondria, occurring at both 5’-and 3’-ends as well as at internal positions ^46^. Especially 3’-editing restores missing nucleotides and base-pairing in the acceptor stem of overlapping tRNA transcripts. For both *R. culicivorax* and *S. magnus*, such events were identified. In *R. culicivorax*, tRNA^Pro^ and tRNA^Val^ overlap by three positions. The overlap remains part of the downstream tRNA^Val^, and the truncated upstream tRNA^Pro^ is completed by the incorporation of three A residues, restoring the acceptor stem and the unpaired position immediately upstream of the CCA-end (discriminator position; Fig. 3, Supp. Table 2A). Furthermore, 3’-end and 3’-connector of tRNA^Tyr^ are heavily edited and carry 11 non-encoded residues that restore a perfectly paired acceptor stem (Fig. 3). Like mt tRNA^Tyr^ of the Japanese land snail *Euhadra herklotsi*, this tRNA does not overlap with adjacent gene sequences, and the processing of the precursor transcript is still a matter of debate ^47^. As predominantly A and C residues are incorporated, poly(A) polymerase and CCA-adding enzyme might represent the responsible editing activities ^48,49^.

In the mt tRNA pool of *S. magnus*, 11 tRNAs exhibit edited 3’-ends, ranging from one to three added A residues (tRNA^Thr^, tRNA^Leu1^, tRNA^Phe^, tRNA^Gln^, tRNA^Ile^, tRNA^Gly^, tRNA^Ser2–2^ and tRNA^Asn^) to more extensive editing including A and C incorporation (tRNA^Arg^, tRNA^Tyr^ and tRNA^Ala/Cys^) (Fig. 5). As editing again leads to the addition of a single-stranded position upstream of the CCA-end as well as acceptor stem mismatches (in tRNA^Tyr^, tRNA^Ala/Cys^), the nucleotide incorporation might also be catalyzed by the template-independent poly(A) polymerase and CCA-adding enzyme. Interestingly, only two pairs of tRNA genes (tRNA^Tyr^ and tRNA^Gln^ as well as tRNA^Ser2–2^ and tRNA^Pro^) share an overlapping region, while the other editing substrates do not. For these non-overlapping tRNAs, the maturation pathway remains unclear (Supp. Table 2B).

### Modification patterns in armless tRNAs

Nucleotide modifications are essential features of tRNAs, influencing structure, stability and interaction with proteins. To the best of our knowledge, no modifications have been identified in armless mt tRNAs so far. Depending on the reverse transcriptase, nucleotide modifications can lead to reverse transcription stops as well as to specific misincorporations, allowing for the identification of modifications by RT-signatures ^50,51^. In addition, we pretreated an aliquot of the tRNA preparation with the demethylase AlkB that specifically removes m^1^A, m^3^C and m^1^G modifications ^23^. The comparative sequence analysis of treated versus untreated RNA preparations identified the methylation patterns in the mt tRNA pools of *R. culicivorax* and *S. magnus*.

In T-armless tRNAs of *R. culicivorax*, a specific RT stop at position A9 appeared in the untreated sample (numbering for conventional tRNAs according to Sprinzl et al.; ^52^), indicating an m^1^A modification, as shown for mt tRNA^Asp^ (Fig. 7A, Supp. Fig. 3). In addition, frequent RT misincorporation was observed at this position, which was sequenced as a U residue in the tRNA, as described for m^1^A ^51^. In the nematode class of Chromadorea, the m^1^A9 modification is essential for the functionality of T-armless tRNAs in terms of aminoacylation rate and EF-Tu binding ^53^, and our data confirm this modification in the corresponding mt tRNAs of *R. culicivorax*. In T-armless mt tRNAs of *S. magnus*, a similar drop in read number is observed at a corresponding position, as shown for tRNA^Gln^ (Fig. 7B, Supp. Fig. 4). However, AlkB treatment did not remove this RT stop, suggesting that the analogous A residue carries a different – not identified - modification. No methylation signals could be detected for D-armless tRNAs of both organisms (Supp. Fig. 3 and 4). For hairpin-like armless tRNAs of *R. culicivorax*, we could not detect any of the AlkB-recognized methylations (Fig. 7A, Supp. Fig. 3). In *S. magnus*, however, armless tRNA^Ile^ and tRNA^Leu1^ show such signals. In the untreated sample of tRNA^Ile^, position C17 shows a drop in coverage upon demethylation treatment, indicating m^3^C at this position (Fig. 7B). For tRNA^Leu2^, U7 and G21 show a slight drop in coverage in the non-treated sample (Supp. Fig. 4). G21 could therefore represent an m^1^G modification. While methylated U is not a typical substrate of AlkB, Clark et al. showed that the corresponding treatment can remove RT misincorporations at such positions ^54^. Hence, the corresponding signal in tRNA^Leu2^ could indicate an m^3^U modification. Besides methylations, RT-signatures allow for the recognition of other modification sites. Within the anticodon loop, bulky modifications stabilize its open conformation^55^. At position A37 of a standard tRNA, these modifications include t^6^A, i^6^A and ms^2^i^6^A (conventional nomenclature used in the literature ^56^) among others that heavily interfere with reverse transcription, leading to a reduction of read numbers and a high misincorporation rate ^51^. We identified such signals for some of the armless mt tRNAs in both organisms (*R. culicivorax*: tRNA^Phe^, tRNA^Ser2^, tRNA^Trp^, tRNA^Leu1^, tRNA^Leu2^, tRNA^Gln^, tRNA^Glu^, tRNA^Thr^; *S. magnus*: tRNA^Phe^, tRNA^Trp^, tRNA^Ser2^, tRNA^Lys^ (Supp. Fig. 3 and 4). In *S. magnus*, we also detected interference with reverse transcription at G37 (standard tRNA numbering) in the anticodon loop, indicating that yW, o^2^yW or m^1^G may be present in tRNA^Pro^, tRNA^His^, tRNA^Leu1^, tRNA^Leu2^, tRNA^Arg^, and tRNA^Ala/Cys^ (Supp. Fig. 4).

**Figure 7:**
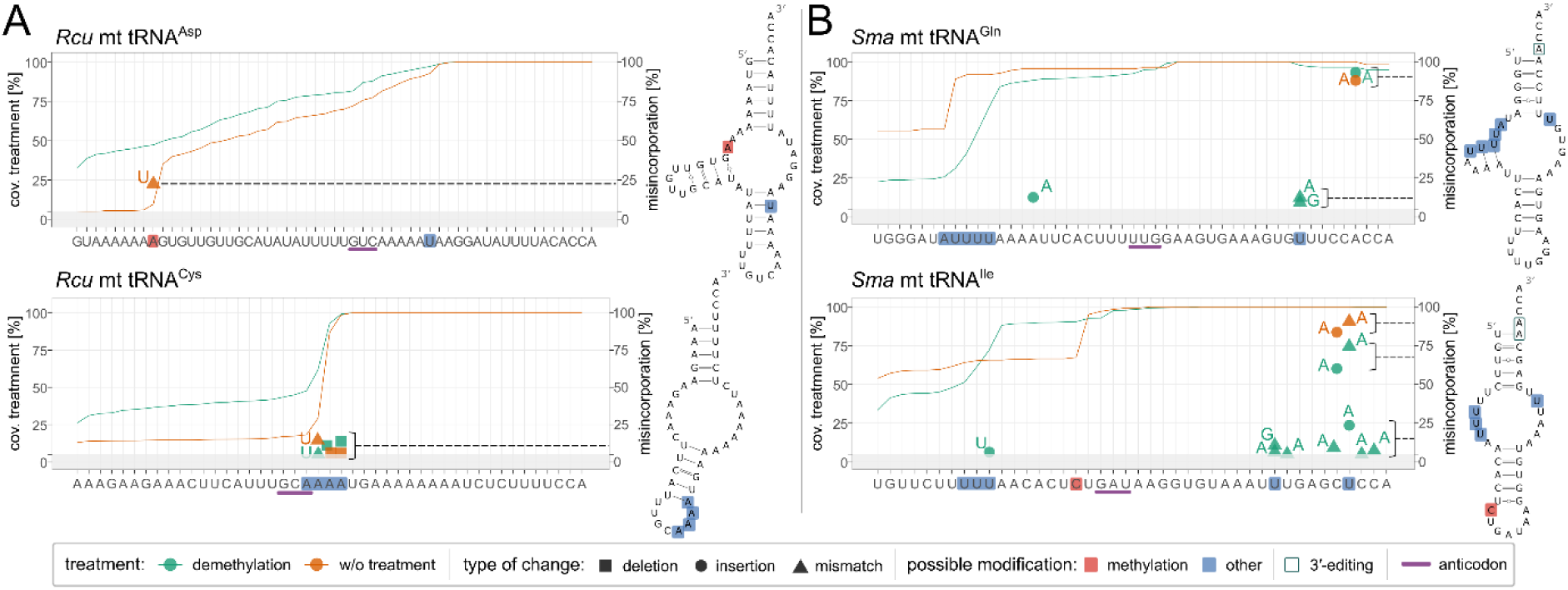
Modification analysis of mt tRNAs. Coverage plots are shown for sequencing data derived from demethylated total RNA (green graph) and non-treated total RNA (orange graph) from *R. culicivorax* (*Rcu*) and *S. magnus* (*Sma*). Starting from the 3’-end, the coverage is normalized to 100% and drops moving towards the 5’-end, a signal characteristic of premature termination of cDNA synthesis. Base modifications lead to misincorporations or to massive RT stops that result in a pronounced drop in sequence reads. Deviations from genomic sequences are highlighted by object forms (deletion, insertion and mismatch), and the corresponding frequency in the reads is indicated by the dashed lines. Putative modification sites are marked in blue, putative methylation sites in red. At certain methylation sites, read drops and misincorporations occur simultaneously, as shown in (**A**). The misincorporation signals in *Rcu* mt tRNA^Cys^ suggest that the 3’-part of the anticodon loop is modified to keep the loop in an open structure. (**B**) In *S. magnus* mt tRNAs for glutamine and isoleucine, single base insertions are found in few read numbers and probably represent the result of polymerase slippage. In *Sma* tRNA^Gln^, an unidentified modification at A7 leads to a signal drop in both demethylated and untreated samples. In both tRNAs, the highly abundant base insertions/deletions close to the tRNA 3’-end represent editing events. To reduce the background signal in the sequence deviations, a threshold of 5% was used (grey bar). The fact that in some cases the number of sequence reads at the 5’-end of the demethylated *S. magnus* mt tRNAs falls below the value for the untreated sample is likely due to a statistical artefact caused by the very small amount of biological material in this analysis. The complete sequence data without threshold are shown in the supplementary data.

Misincorporations not related to tRNA modification are found at the 3’-end of several tRNAs. These sequence aberrations are the result of posttranscriptional editing events in both *S. magnus* and *R. culicivorax* (Fig. 7B, Supp. Fig. 3 and 4). In *S. magnus*, some misincorporations are located in the connector loop of armless tRNAs. Whether these signals indicate modifications keeping the loop regions in a single-stranded conformation or represent misincorporations due to reverse transcriptase slippage in homopolymeric sequence repeats remains to be clarified.

## Conclusions

The characterized mitochondrial tRNAs display strong deviations from their canonical counterparts. With an average size of 45 nt (*S. magnus*) to 52 nt (*R. culicivorax*), they are highly reduced compared to the standard size of 76 nt. Yet, our data show that the miniaturized mt tRNAs are expressed and processed by a series of maturation activities like CCA-adding enzyme, the tRNA editing machinery as well as modification enzymes. In Enoplea, miniaturized and armless mt tRNAs are highly conserved, and tRNAs of the same identity exhibit the same structure throughout the different organisms of this class.

The 3’-editing events in several mt tRNAs of both species remove mismatches in the predicted sequences and stabilize the acceptor stem, comparable to the known cases in snails and other organisms ^57^. The fact that genome-based predictions indicate mismatches at these (unedited) positions underscores the importance of transcriptome analysis to identify the actual tRNA sequences, and, consequently, their functionality. In the mt pool of *S. magnus*, this becomes even clearer, as the confirmed tRNA sequences and gene locations vary greatly from the predictions described in several studies ^5,14–18^.

The mt tRNA set of *T. urticae*, related to *S. magnus* with a phylogenetic location in the same superorder of Acariformes, also shows miniaturization and strong deviations from canonical tRNAs ^21^. In contrast to the conserved tRNA sequences and structures in Enoplea mitochondria ^6^, mite mt tRNAs with the same identity differ dramatically in these features (Table 1). This indicates that the loss of tRNA arms occurred multiple times independently in evolution.

**Table 1:**
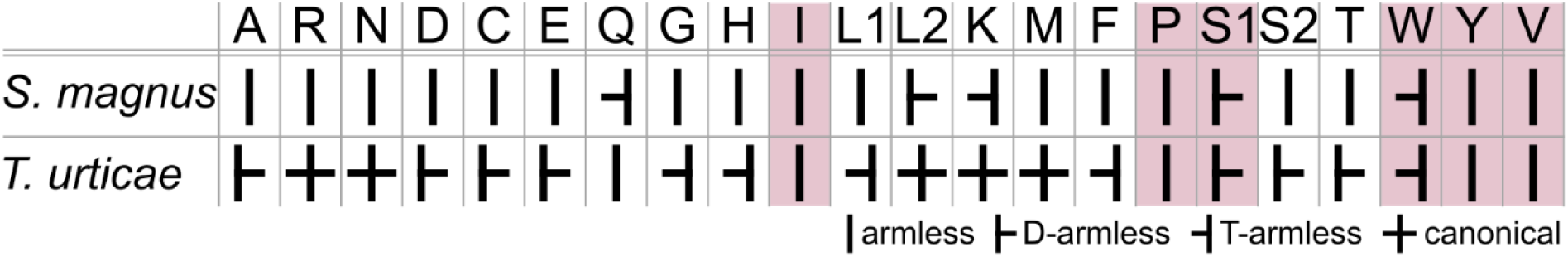
Comparison of the confirmed mitochondrial tRNAs depicted by their structure in the two Acariformes *S. magnus* and *T. urticae*.

In canonical tRNAs, the T-and D-arms play a key role in the formation of the tertiary structure, shaping the elbow region that is crucial for the interaction with proteins ^58^. Since these elements are absent in armless mt tRNAs, the question arises as to how these tRNAs can still interact with proteins. In functional studies, it was shown that the CCA-adding enzyme as well as the mitochondrial elongation factor EF-Tu1 of *R. culicivorax* are highly adapted to their armless tRNA substrates ^37,38^. Obviously, the mitochondrial translation machinery exhibits a surprisingly high evolutionary plasticity that compensates for the structural reductions of its tRNAs. Further experiments are required to understand the functional interplay of the individual components of this unusual translation system.

## Methods

### Specimen collection and RNA isolation

Specimen of *R*. *culicivorax* were cultivated as described ^59^, in accordance with a protocol established by Edward Platzer (UC Riverside, USA, personal communication), who also provided the nematode source material. Specimen of *S*. *magnus* were collected in the Göttinger Wald (Lower Saxony) and separated from soil using heat extraction ^60^ and collected alive in water. Individuals were identified under a stereomicroscope using a determination key ^61^, collected in 96% ethanol and stored at -20°C until use. The samples of both species were homogenized as follows: 100 mg of *R. culicivorax* were transferred to a tube of the Precellys^®^ Lysing Kit Soft tissue homogenizer CK14 (bertin technologies). 1 ml TRIzol^TM^ (Thermo Fisher Scientific) was added, and the material was homogenized twice for 30 s at 6 m/s in a FastPrep-24 homogenizer. 37 mg of *S. magnus* were frozen in liquid nitrogen and ground in a cellcrusher tissue pulverizer (Cellcrusher) according to the manufacturer’s instructions. The material was dissolved in 750 µl TRIzol^TM^ and homogenized for 30 s at 6 m/s in a FastPrep-24 homogenizer. Phase separation of both samples was performed according to the standard protocol for TRIzol^TM^ extraction (Thermo Fisher Scientific). The obtained aqueous phase was purified using the Monarch^®^ Spin RNA Cleanup Kit (New England Biolabs) according to the RNA purification protocol. The quality of the total RNA preparations was checked on a 1 % agarose gel.

### Preparation of recombinant AlkB

The plasmid pET-22b(+), containing the sequence for the *E. coli* DH5α demethylase (AlkB) with a C-terminal His-tag, was used for recombinant expression in *E. coli* BL21(DE3) cells. Two 400 ml LB cultures with 100 µg/ml ampicillin were grown at 37 °C until an OD600 of 0.6. Expression was induced by the addition of IPTG and FeSO_4_ to a final concentration of 1 mM and 5 µM, respectively, and further incubation for an additional 4 h at 30 °C. Cells were harvested by centrifugation and used for protein purification with buffers based on the protocol of Wang et al. ^62^. The cell pellet was resuspended in lysis buffer (10 mM Tris/HCl pH 7.4, 300 mM NaCl, 10 mM MgCl_2_, 2 mM CaCl_2_, 2 mM 2-Mercaptoethanol, 5 % Glycerol) and disrupted using Zirconium beads (Fisher Scientific) and the FastPrep-24 homogenizer for 30 s at 6 m/s. The suspension was centrifuged, and the obtained supernatant was sterile-filtered (TPP^®^). The filtrate was applied to a HisTrap^TM^ HP column (Cytiva) and purified using the ÄKTA pure^TM^ chromatography system (Cytiva). The protein was eluted with 10 mM Tris/HCl pH 7.4, 200 mM NaCl_2_, 10 mM 2-Mercaptoethanol, 300 mM imidazole. Protein fractions (determined by absorption values at 280 nm) were diluted with buffer A (10 mM Tris/HCl pH 7.4, 10 mM 2-Mercaptoethanol) and applied to a HiTrap^TM^ Heparin HP column (Cytiva). The elution was performed by a linear gradient of buffer B (10 mM Tris/HCl pH 7.4, 10 mM 2-Mercaptoethanol, 1 M NaCl). The buffer of the protein fractions was exchanged by the storage buffer (10 mM Tris/HCl pH 7.4, 250 mM NaCl) using a Vivaspin 20 column (MWCO 10 kDa; Sartorius). Glycerol was added to a final concentration of 30% and the protein was stored at -70 °C.

### ARM-seq and LOTTE-Seq

Total RNA was treated with AlkB demethylase according to the published protocol ^23^ with minor changes: The volume of the reaction mixture was adjusted to 50 µl containing 2 µg of total RNA and 160 pmol of recombinant AlkB. The reaction was stopped by adding 10 mM EDTA, and the RNA was purified using the Monarch^®^ Spin RNA Cleanup Kit (New England Biolabs).

Specific tRNA ligation was performed using an advanced hairpin adapter (5’-pCCGCGGCCGBBBBBBBBTGGAATTCTCGGGTGCCGGGAGAAGGCACCCGAGAATTCCATTTTTTTTCGGCCGCGGTGGNp-3’; B=T,C,G) compared to (Erber et al. 2020) (Supp. Table 1). The reaction mixture contained 200 pmol hairpin adapter, 2 µg of untreated or demethylated total RNA, 200 U T4 DNA ligase (New England Biolabs), 66 mM Tris/HCl pH 7.6, 6.6 mM MgCl_2_, 10 mM DTT, 66 µM ATP and 25% (v/v) DMSO. The reaction was incubated for 8 h at 32 °C and stopped by heat inactivation of the enzyme at 65 °C for 10 min. The ligated RNA was purified using the Monarch^®^ Spin RNA Cleanup Kit (New England Biolabs).

90 pmol of RT primer (spiked with 10 pmol 5’-^32^P-labeled RT primer) were added to the ligation product. After the addition of NTPs to a final concentration of 0.5 mM dATP/dGTP/dCTP, 0.35 mM dTTP and 0.15 mM Biotin-16-dUTP in a total volume of 14 µl, the mixture was incubated for 5 min at 65° C. After a cooling step on ice for 1 min, 1x first strand buffer, 5 mM DTT and 200 U SuperScript^TM^ III were added to a total volume of 20 µl. After incubation at 55 °C for 30 min, the RNA was degraded by adding 250 mM NaOH and heating the sample for 3 min to 95 °C. RNA degradation was stopped by adding 250 mM HCl. cDNA was purified by denaturing PAGE (15 %). cDNA bands of expected length were cut from the gel and eluted in 10 mM Tris/HCl pH 7.5, 200 mM NaCl and 1 mM EDTA. The cDNA was ethanol-precipitated in the presence of 10 µg carrier LPA (Thermo Fisher Scientific).

The cDNA was circularized using the TS2126 RNL and purified using hydrophilic streptavidin magnetic beads (New England Biolabs) as described ^24^. The circularized cDNA was PCR-amplified with primers that introduced flow cell linkers and indices. A 50 µl reaction contained 5 µl of cDNA,1 U Phusion DNA polymerase (Thermo Fisher Scientific), 1x Phusion HF buffer, 0.2 mM dNTPs, 0.5 µM of Illumina PCR and respective Index primer (Supp. Table 1). The cycling program was performed as follows: 98 °C for 30 s; [98 °C for 10 s; 60 °C for 20 s; 72 °C for 15 s] 18x; 72 °C for 2 min. The PCR product was purified on an 8 % native PAGE. DNA with a size above 140 bp was eluted in 10 mM Tris/HCl pH 7.5, 200 mM NaCl and 1 mM EDTA, followed by ethanol precipitation in the presence of 5 µg carrier LPA (Thermo Fisher Scientific). Purified libraries were used for Illumina sequencing on a NovaSeq 6000 platform (configuration: 2x 150 bp) at Genewiz^®^ Germany GmbH (Leipzig). A list of all used oligonucleotides is provided in Supp. Table 1.

### Bioinformatical analysis of LOTTE-Seq data

Data analysis of the obtained sequencing data (total tRNA pools) was partly conducted following the best-practice workflow ^63^, with adjustments for mitochondrial tRNA search.

Artificial genomes were constructed in several steps. Nuclear-encoded mitochondrial genome parts were identified with exonerate (ver. 2.0.0) ^64^ (minimum score 500) in nuclear genome and hard-masked with bedtools (ver. 2.27.1) ^65^. If tRNA annotations were available (for *R. culicivorax*), artificial chromosomes made from mt-tRNA sequences, and mt tRNAs were hard-masked in the mitochondrial genome. The structure of artificial chromosomes includes 3 bases from the upstream tRNA region, as well as an additional CCACCA-end that allowed for the detection of tRNA molecules that had been marked for degradation but were still present in the mt tRNA pool ^66^. Redundant mt tRNA sequences were collapsed. Reference genome GCA_963922205.1 and mitogenome NC_008640.1 were used for *R. culicivorax* mt tRNA analysis. Annotation of mt tRNAs was obtained with species-specific CMs ^11^. Because mt tRNA annotations in *S. magnus* are inconsistent, the first round of alignment was performed using a non-masked mitochondrial genome to localize the tRNAs. The next round included masked tRNAs as described for *R. culicivorax*. Reference data used for *S. magnus* analysis: GCA_034698365.1 genome and NC_011574.1 mitogenome.

Sequencing data processing included quality control with FastQC ^67^, followed by adapter trimming with cutadapt (ver. 4.7) ^68^ and UMI deduplication with umi_tools (ver. 1.1.5) ^69^. Only reads with a length of more than 8 nt and having a CCA-end in sequence were saved for alignment.

Alignment was performed using segemehl (ver. 0.3.4) ^70^ with the following parameters: evalue 500, differences 3, maxinterval 1000, and accuracy 80. Relaxed parameters help to detect reads with mismatches connected to modifications. Visualization was done with the IGV browser ^71^ and RStudio. Genomic region visualization was performed using the circlize package (ver. 0.4.16) ^72^.

Modification profiles were generated from SAM-formatted alignments produced with segemehl. Coverage and sequence differences relative to the reference sequence were derived using CIGAR information. Coverage was normalized to the maximum coverage of each molecule, and modification frequencies were calculated as the percentage of reads supporting a given modification relative to the coverage at each base position. Insertions are indicated at the position of the upstream base and should be interpreted as occurring between the marked base and the following base. tRNA modification profile of *R. culicivorax*: filters were applied to display only coverage and sequence changes supported by at least 50 reads. tRNA modification profile of *S. magnus*: adjusted thresholds were used for lower-coverage molecules (e.g., 5 reads for tRNA^Val^ and 30 reads for tRNA^Glu^, tRNA^Thr^, tRNA^Leu1^), while a threshold of 50 reads was retained for all other molecules.

### Led-Seq

For *ex vivo* probing of native mt tRNAs, total RNA of *R. culicivorax* was depleted of rRNA by adding 0.5 M NaCl and 5% PEG8000. The solution was incubated for 30 min at -20 °C and centrifuged for 30 min at 4 °C and 10,000 rcf. The supernatant was removed, and the procedure repeated. The final solution was precipitated with ethanol. 400 ng of the RNA preparation was heated to 90 °C for 1 min, followed by a refolding step at RT for 5 min. The sample was incubated for 10 min at 37° C in 50 mM Tris/HCl (pH7.5), 100 mM NH_4_Cl, 10 mM MgCl_2_ and 1 mM Pb(OAc)_2_. The reaction was stopped by adding 20 mM EDTA and the RNA was purified using the Monarch^®^ Spin RNA Cleanup Kit (New England Biolabs). The lead-treated small RNA fraction was used for subsequent library preparation and sequenced on an Illumina NovaSeq 6000 platform (configuration: 2x 150 bp) at Genewiz^®^ Germany GmbH (Leipzig) as described ^24^.

The derived data were processed ^24^, and the normalized probing signal *S* was evaluated and mapped to the corresponding structure. Additionally, a moving average was applied to *S* using a Savitzky-Golay method (convolution coefficient for smoothing with a window size of 5) ^73^:

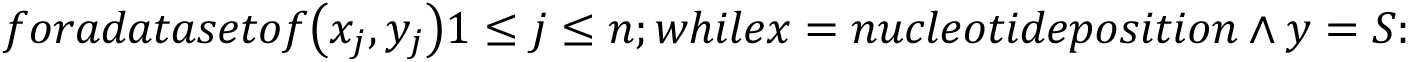

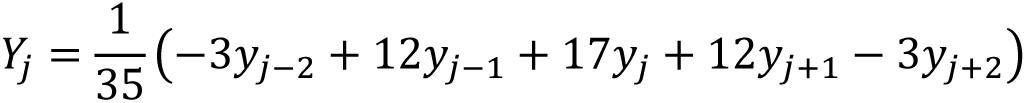

Resulting negative values for Y_j_ were set to zero.

### In-line probing of *in vitro* transcribed tRNAs

In-line probing of *in vitro* transcribed mt tRNA^Ala/Cys^ and mt tRNA^Cys/Ala^ was performed as described ^25^.

## Data availability

The oligonucleotides used in this study for Illumina library preparation are included in the supplementary table 1. The sequencing data are available in the NCBI Sequence Read Archive and connected with BioProject ID PRJNA1477637. A computational pipeline is accessible at: https://github.com/julie-tooi/aberrant_mttrnas.

## Supplementary Data

Supplementary Data are available online.

## Acknowledgements

We thank Stefan Scheu and Alexander Brandt for providing specimens and mitochondrial genome data of *Steganacarus magnus*. We also thank Tobias Friedrich for technical assistance and Lowell George for title inspiration.

## Author Contributions

PFS and MM conceived the study. PFS, HB and MM supervised and evaluated the work. JG performed all biochemical and sequencing experiments and optimized the LOTTE-Seq approach. TK and SvL performed Led-Seq analysis, and IO analyzed and evaluated the LOTTE-Seq data. PS and IS raised and collected specimen of *R. culicivorax* and *S. magnus*, respectively. All authors wrote, edited and approved the manuscript.

