## Supplementary Material for "A little feat in translation: miniaturized tRNAs in nematode and arachnid mitochondria"

### Supplementary figures

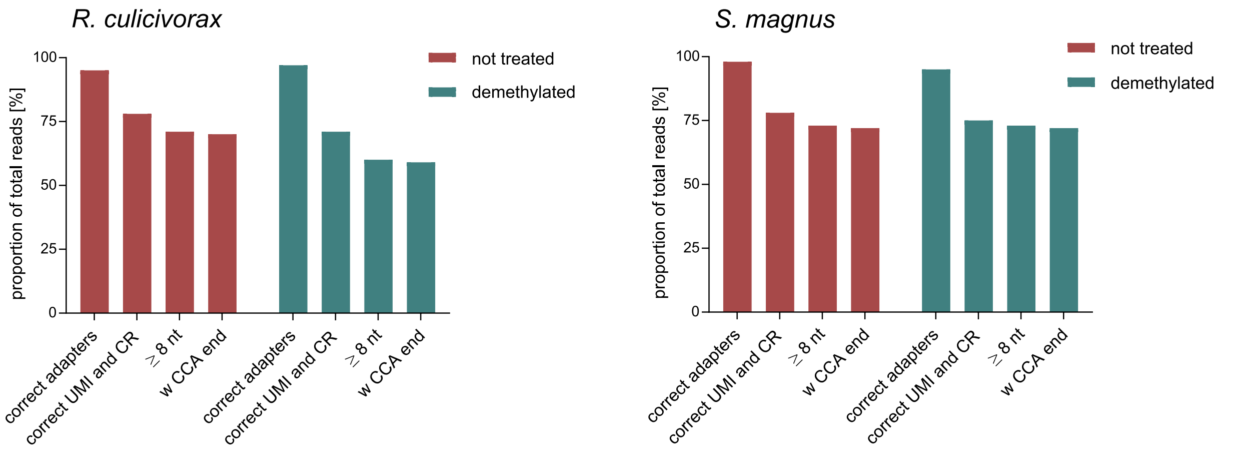


Supplementary figure 1: **Quality of sequencing libraries from *R. culicivorax* and *S. magnus***

The bar graphs show read processing statistics after Illumina sequencing of libraries derived from untreated total RNA (red) and libraries derived from demethylated total RNA (green). Bars represent the proportion of reads compared to the total amount of reads in processing order from left to right: percentage of reads that carry both Illumina adapters (correct adapters); percentage that also carry the UMI sequence and constant region from the hairpin adapter (correct UMI and CR); percentage that are longer than 8 nt after trimming (≥ 8 nt); percentage that carry also a 3′-CCA-end (w CCA end). Trimmed reads with a minimum size of 8 nt and a 3′-CCA-end were used for subsequent alignment.

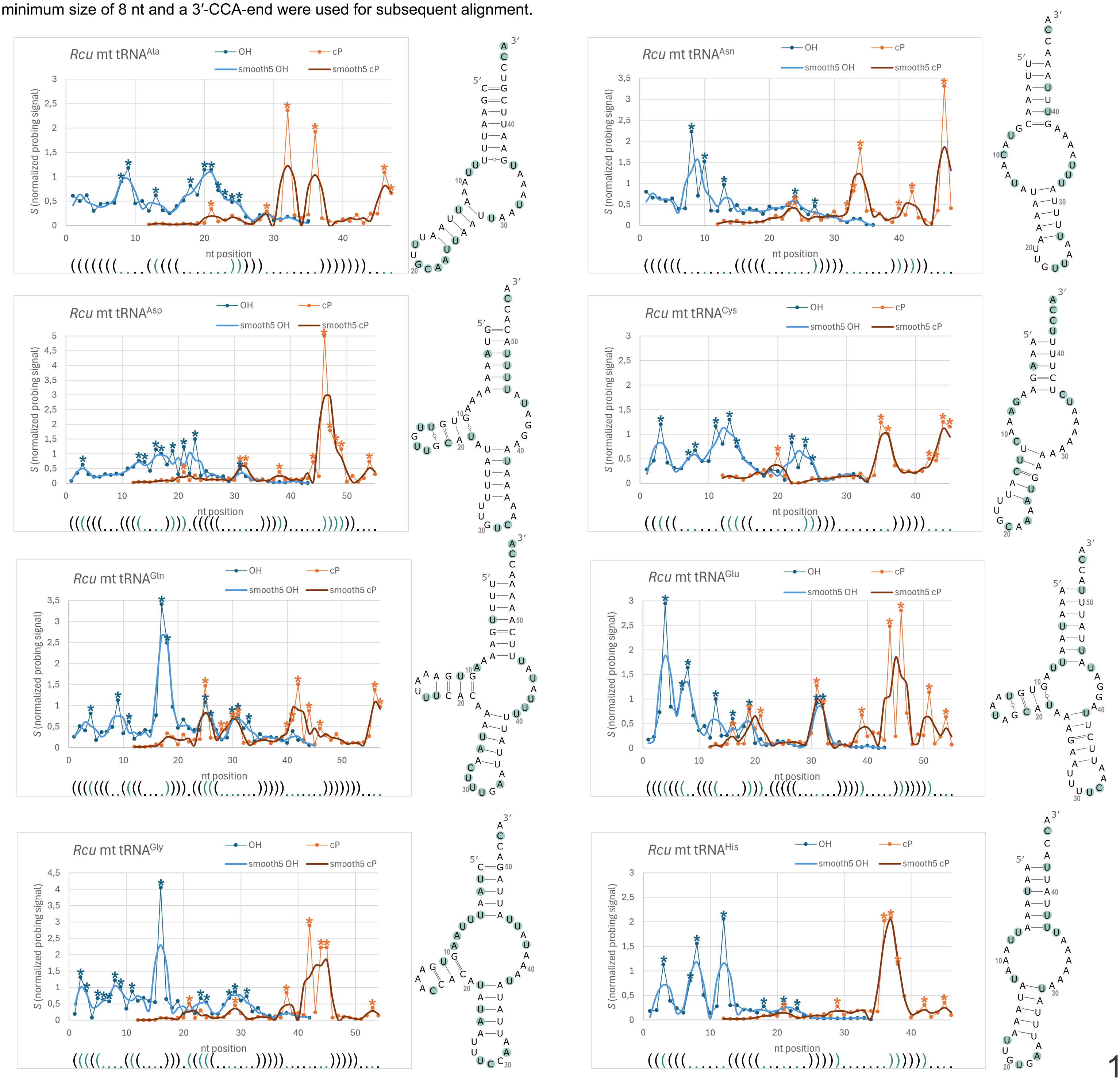

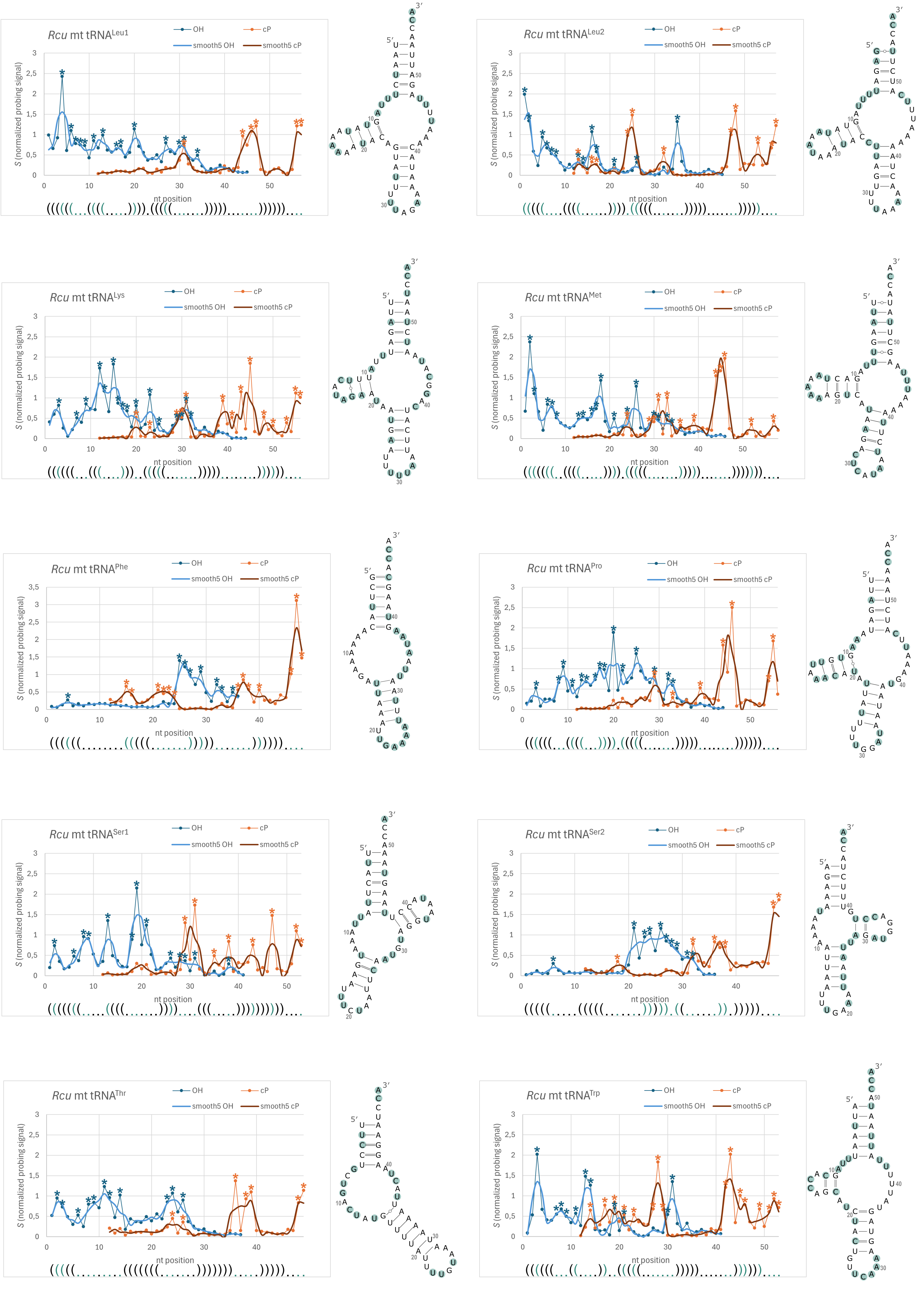
Rcu mt tRNALeu1
Rcu mt tRNALeu2
Rcu mt tRNALys
Rcu mt tRNAMet
Rcu mt tRNAPhe
Rcu mt tRNAPro
Rcu mt tRNASer1
Rcu mt tRNASer2
Rcu mt tRNAThr
Rcu mt tRNATrp
OH
cP
smooth5 OH
smooth5 cP
S (normalized probing signal)
nt position

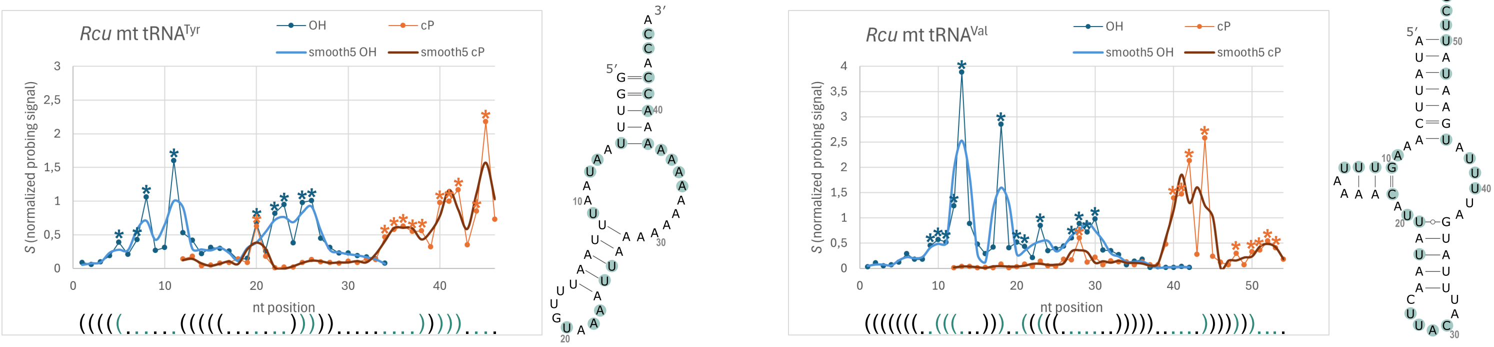


Supplementary figure 2: **Led-seq probing signal of mt tRNAs from *R. culicivorax***

Shown are the graphs for each tRNA depicting the normalized probing signal *S* (Y-axis) at each nucleotide position (X-axis) for both libraries (cyclophosphat-libraries: cP; OH-libraries: OH). The intensity of the signal correlates directly with cleavage at this position. Peaks marked with * show a high signal intensity compared to their neighbouring signals and are therefore more likely being unpaired or are effected by fraying. Next to the graph the corresponding tRNA is depicted, where green marked nucleotides represent high intensity peaks marked with *. In addition a moving average was applied to the probing signal with a window size of 5 (Savitzky-Golay method) for both libraries (smooth5 OH and smooth5 cP). The dot bracket notation is aligned to the signal for each graph.

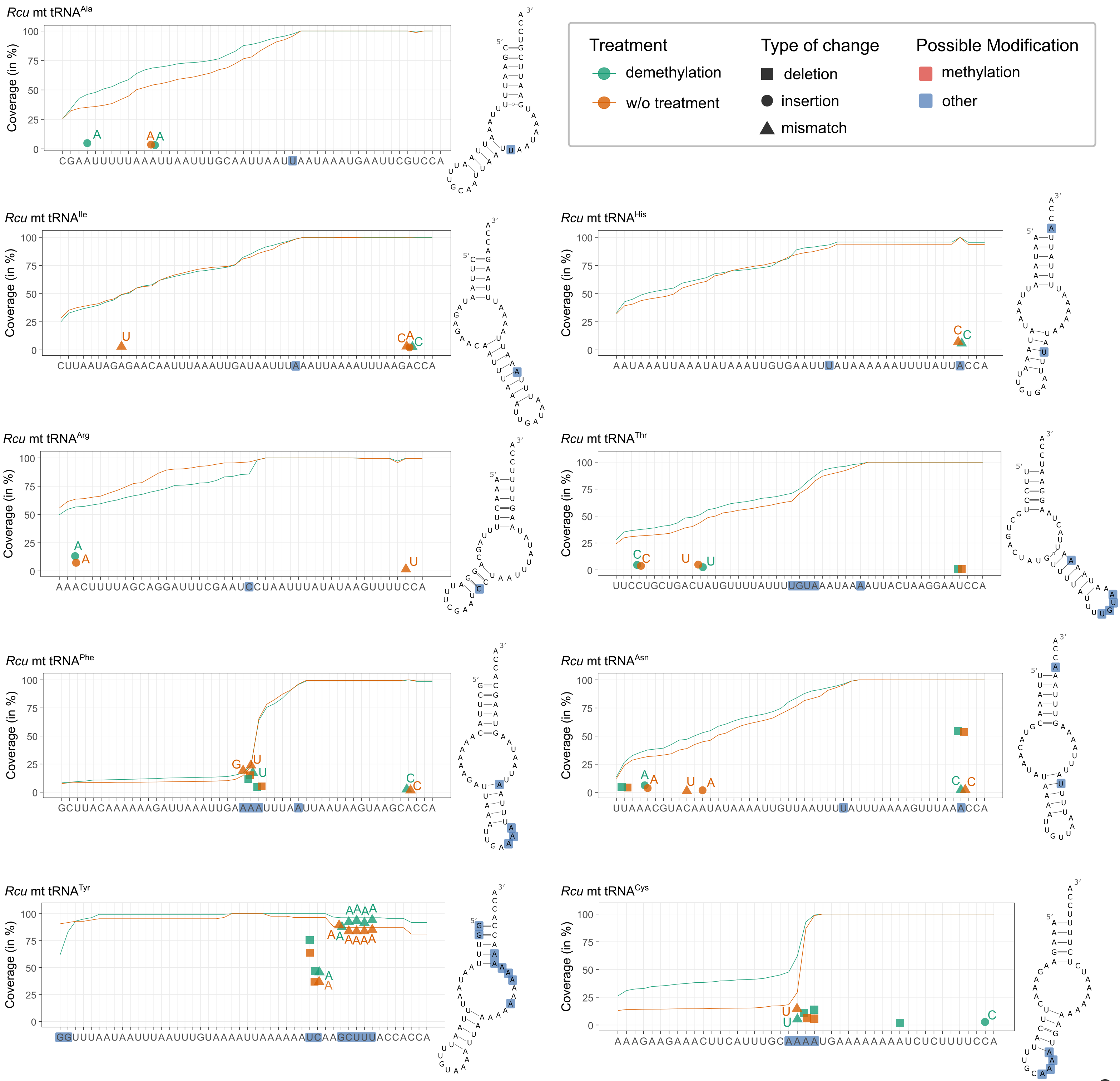

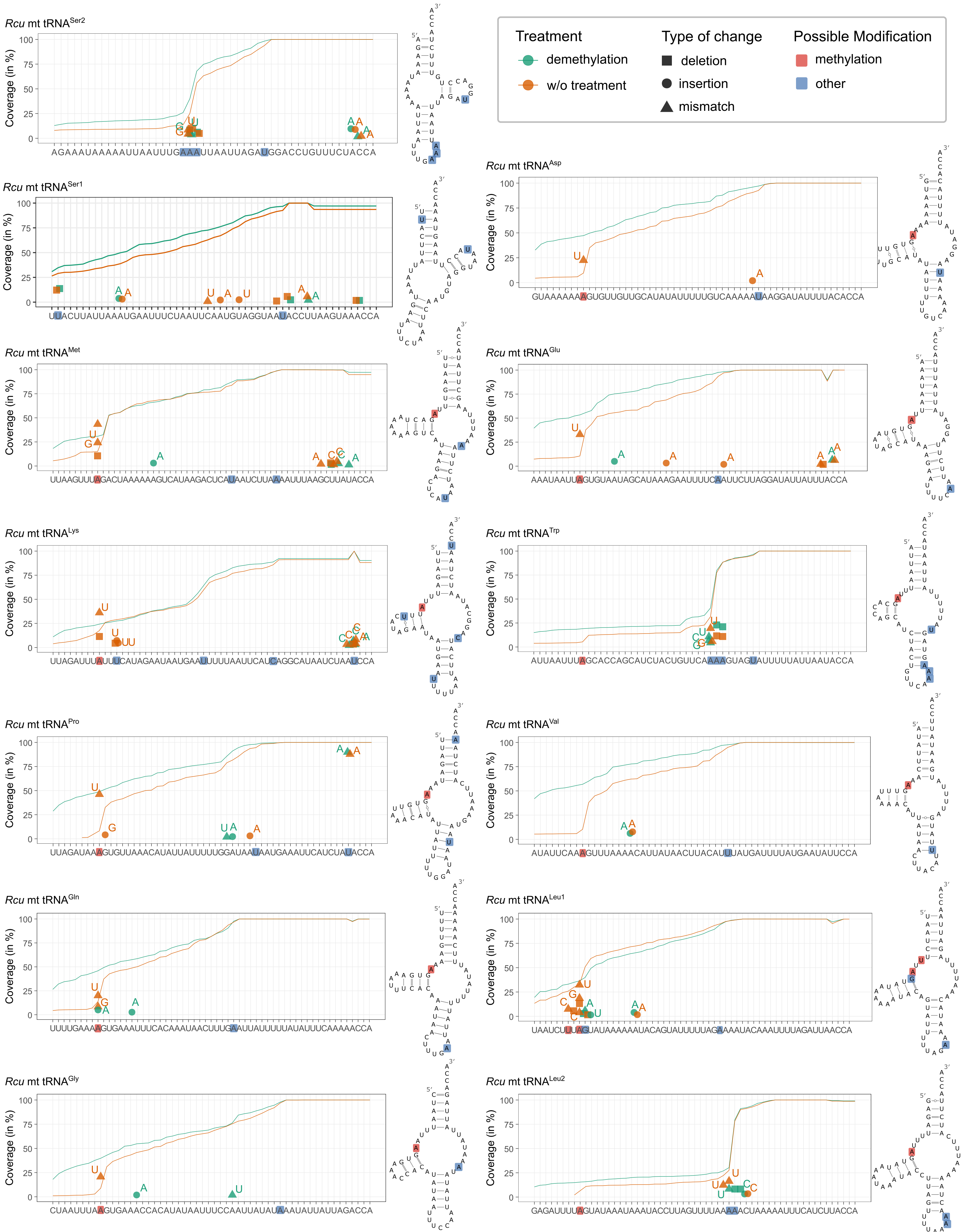


Supplementary figure 3: **Modification analysis of mt tRNAs from *R. culicivorax***

Shown are the coverage plots for each tRNA depicting the coverage along the whole tRNA sequence for demethylated and non-demethylated libraries (green and orange curve, respectively). All types of changes within the reads compared to the sequence are marked in the same plots. High mismatch areas are marked as well as pronounced coverage drops are marked in the sequence (blue and red nucleotides), where red-marked positions can be traced back to methylated nucleotides and blue-marked positions could be due to other modifications or structure dependent RT stops. Notably percentage of coverage and misincorporation were counted seperately and are depicted in one graph for representation (see figure 7).

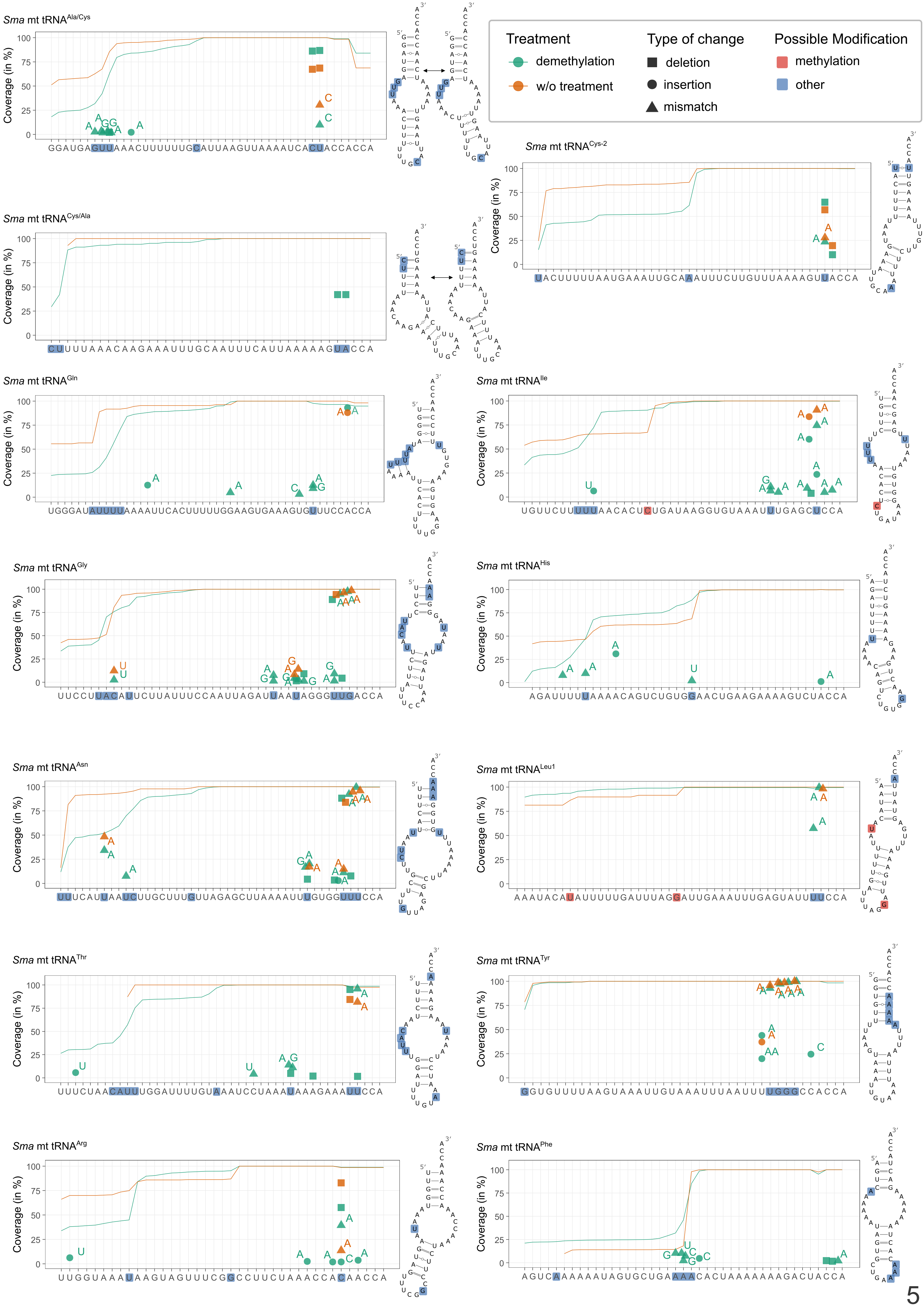

Treatment
demethylation
w/o treatment
Type of change
deletion
insertion
mismatch
Possible Modification
methylation
other
Sma mt tRNA Ala/Cys
Sma mt tRNA Cys-2
Sma mt tRNA Cys/Ala
Sma mt tRNA Gln
Sma mt tRNA Ile
Sma mt tRNA Gly
Sma mt tRNA His
Sma mt tRNA Asn
Sma mt tRNA Leu1
Sma mt tRNA Thr
Sma mt tRNA Tyr
Sma mt tRNA Arg
Sma mt tRNA Phe
Coverage (in %)

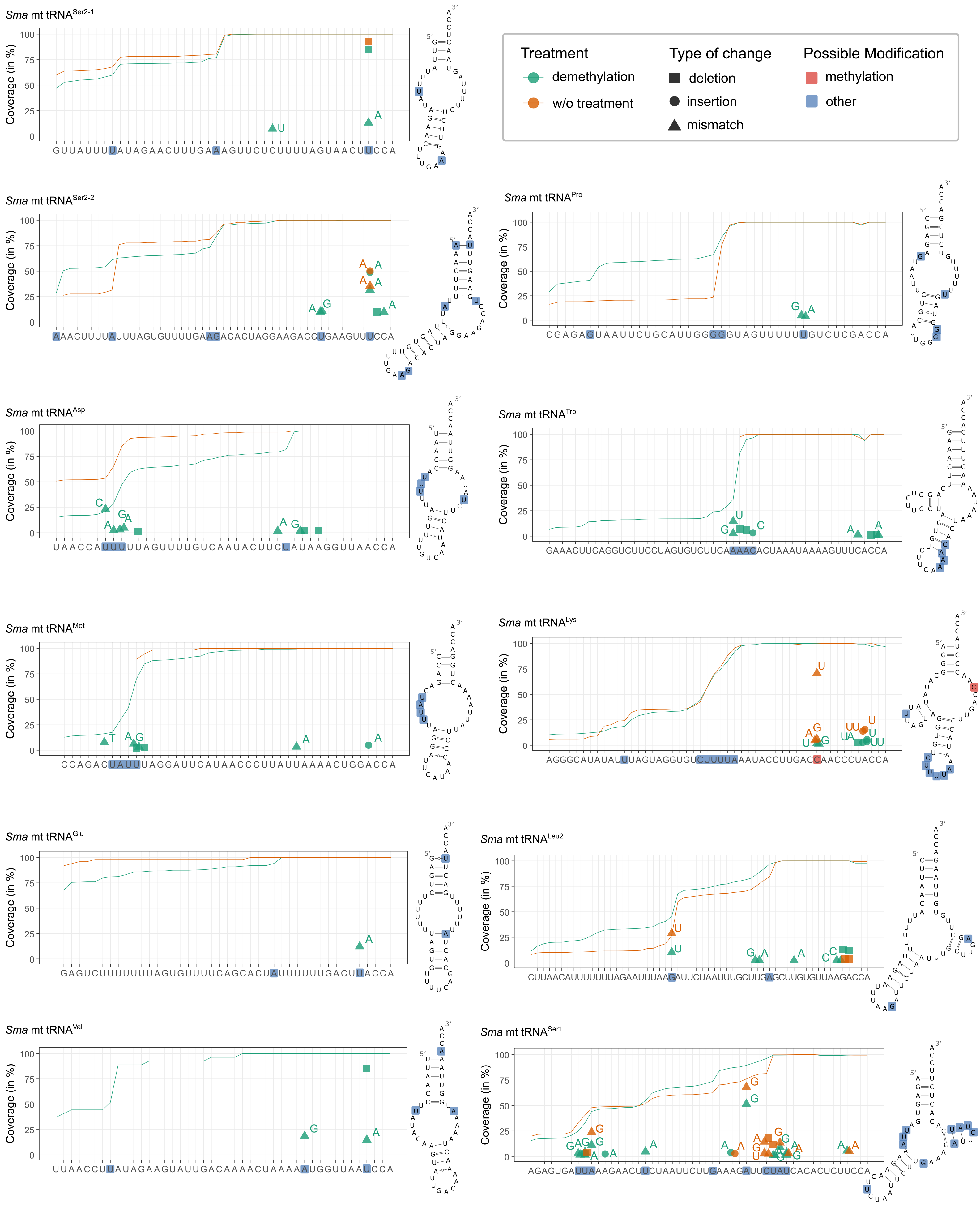


Supplementary figure 4: **Modification analysis of mt tRNAs from *S. magnus***

Shown are the coverage plots for each tRNA depicting the coverage along the whole tRNA sequence for demethylated and non-demethylated libraries (green and orange curve, respectively). All types of changes within the reads compared to the sequence are marked in the same plots. High mismatch areas are marked as well as pronounced coverage drops are marked in the sequence (blue and red nucleotides), where red-marked positions can be traced back to methylated nucleotides and blue-marked positions could be due to other modifications or structure dependent RT stops. Notably percentage of coverage and misincorporation were counted seperately and are depicted in one graph for representation (see figure 7).

**Supplementary Table 1: List of Oligonucleotides for LOTTE-seq/ARM-seq**

| # | oligo description | sequence | modifications | comments | company |
|---|---|---|---|---|---|
| 1 | LOTTE-seq hairpin adapter | 5’-pCCGCGGCCGBBBBBBBBTGGAATTCTCGGGTGCCGGGAGAAGGCACCCGAGAATTCCATTTTTTTTCGGCCGCGGTGGNp-3’ | 5’- and 3’-phosphorylation | B=T,C,G; N=A,T,C,G | Eurofins Genomics |
| 2 | RT primer circularization | 5’-GATCGTCGGACTGTAGAACTCTGAAC/HEG/CACTCA/HEG/GCCTTGGCACCCGAGAATTCCA-3’ | 5’- phosphorylation; two internal HEG mod | HEG= hexaethylen glycol | Microsynth AG |
| 3 | forward Primer PCR | 5’-AATGATACGGCGACCACCGAGATCTACACGTTCAGAGTTCTACAGTCCGA-3’ | *none* | *none* | Microsynth AG |
| 4 | reverse Primer PCR -Index 1 | 5’-CAAGCAGAAGACGGCATACGAGATCGTGATGTGACTGGAGTTCCTTGGCACCCGAGAATTCCA-3’ | *none* | *none* | Microsynth AG |
| 5 | reverse Primer PCR -Index 2 | 5’-CAAGCAGAAGACGGCATACGAGATACATCGGTGACTGGAGTTCCTTGGCACCCGAGAATTCCA-3’ | *none* | *none* | Microsynth AG |
| 6 | reverse Primer PCR -Index 3 | 5’-CAAGCAGAAGACGGCATACGAGATGCCTAAGTGACTGGAGTTCCTTGGCACCCGAGAATTCCA-3’ | *none* | *none* | Microsynth AG |
| 7 | reverse Primer PCR -Index 4 | 5’-CAAGCAGAAGACGGCATACGAGATTGGTCAGTGACTGGAGTTCCTTGGCACCCGAGAATTCCA-3’ | *none* | *none* | Microsynth AG |

**Supplementary Table 2: List of sequenced mt tRNAs with their genome location and coverage**

comment: nucleotides represented in grey are posttranscriptionally added/edited nucleotides and do not show in mt genome sequence

**Table 2A: *Romanomermis culicivorax***

| | | | coverage unique filter | | coverage of reverse oriented reads | |
|---|---|---|---|---|---|---|
| **tRNA** | **sequence in 5' -3' direction** | **mt genome position (NCBI reference NC_008640.1)** | **demethylated** | **untreated** | **demethylated** | **untreated** |
| Ala | CGAATTTTTAAATTAATTTGCAATTAATTAATAAATGAATTCGTCCA | 3986..4029 (+) | 3785 | 3833 | 415 | 403 |
| Cys | AAAGAAGAAACTTCATTTGCAAAATGAAAAAAAATCTCTTTTCCA | 14881..14922 (-) | 3278 | 2254 | 49 | 100 |
| Asp | GTAAAAAAAGTGTTGTTGCATATATTTTTGTCAAAAATAAGGATATTTTACACCA | 17397..17448 (+) | 2661 | 3418 | 3 | 7 |
| Glu | AAATAATTAGTGTAATAGCATAAAGAATTTTCAATTCTTAGGATATTATTTACCA | 4183..4234 (-) | 2229 | 3085 | 3 | 5 |
| Phe | GCTTACAAAAAAGATTAAATTGAAAATTTAATTAATAAGTAAGCACCA | 17513..17557 (+) | 3902 | 4031 | 528 | 582 |
| Gly | CTAATTTAAGTGAAACCACATATAATTTCCAATTATATAAATATTATTAGACCA | 17458..17508 (+) | 8874 | 6935 | 10 | 9 |
| His | AATAAATTAAATATAAATTGTGAATTTATAAAAAAATTTTATTACCA | 12825..12867 (+); 24829..24871 (-) | 4512 | 4540 | 2 | 1 |
| Ile | CTTAATAGAGAACAATTTAAATTGATAATTTAAATTAAAATTTAAGACCA | 3214..3260 (+) | 6763 | 4473 | 1 | 2 |
| Lys | TTAGATTTATTTCATAGAATAATGAATTTTTAATTCATCAGGCATAATCTAATCCA | 8258..8310 (-); 11290..11342 (-); 22189..22241 (+) | 15688 | 17864 | 85 | 133 |
| Leu1 | TAATCTTTAGTATAAAAAATACAGTATTTTTAGAAAATACAAATTTTAGATTAACCA | 3920..3973 (-) | 8066 | 7806 | 6 | 6 |
| Leu2 | GAGATTTTAGTATAAATAAATACCTTAGTTTTAAAAACTAAAAATTTCATCTTACCA | 1514..1567 (+) | 2140 | 2447 | 15 | 24 |
| Met | TTAAGTTTAGACTAAAAAAGTCATAAGACTCATAATCTTAAAATTTAAGCTTATACCA | 17336..17390 (+) | 4268 | 3774 | 7 | 6 |
| Asn | TTAAACGTACAATATAAAATTGTTAATTTTATTTAAAAGTTTAAACCA | 6915..6959 (-); 9947..9991 (-); 20517..20561 (+); 23540..23584 (+) | 11117 | 13073 | 158 | 165 |
| Gln | TTTTGAAAAGTGAAATTTCACAAATAACTTTGAATTATTTTTATATTTCAAAAACCA | 1032..1085 (-) | 5523 | 5208 | 0 | 0 |
| Arg | AAACTTTTAGCAGGATTTCGAATCCTAATTTATATAAGTTTTCCA | 3161..3202 (-) | 3381 | 3676 | 7 | 20 |
| Ser1 | TTACTTATTAAATGAATTTCTAATTCAATGTAGGTAATACCTTAAGTAAACCA | 4040..4089 (+) | 7795 | 9892 | 27 | 36 |
| Ser2 | AGAAATAAAAATTAATTTGAAATTAATTAGATGGACCTGTTTCTACCA | 7846..7890 (+); 10878..10922 (+); 22609..22653 (-) | 6215 | 6096 | 770 | 778 |
| Thr | TTCCTGCTGACTATGTTTTATTTTGTAAATAAAATTACTAAGGAATCCA | 12906..12951 (+); 24745..24790 (-) | 5786 | 6491 | 16 | 19 |
| Val | ATATTCAAAGTTTAAAACATTATAACTTACATTTATGATTTTATGAATATTCCA | 718..768 (+) | 3190 | 3112 | 3 | 4 |
| Trp | ATTAATTTAGCACCAGCATCTACTGTTCAAAAGTAGTATTTTTATTAATACCA | 26139..26188 (+) | 3433 | 3102 | 0 | 0 |
| Tyr | GGTTTAATAATTTAATTTGTAAAATTAAAAAAAAAAAAAACCACCA | 14944..14975 (+) | 160 | 85 | 0 | 4 |
| Pro | TTAGATAAAGTGTTAAACATATTATTTTTGGATAATAATGAAATTCATCTAAACCA | 668..720 (+) | 3918 | 4247 | 0 | 0 |

**Table 2B:** ***Stegenacarus magnus***

| | | | coverage unique filter | | coverage of reverse oriented reads | |
|---|---|---|---|---|---|---|
| **tRNA** | **sequence in 5' -3' direction** | **mt genome position (NCBI reference NC_011574.1)** | **demethylated** | **untreated** | **demethylated** | **untreated** |
| Ala/Cys | GGATGAGTTAAACTTTTTTGCATTAAGTTAAAATCAACCACCA | 4162..4197 (+) | 5994 | 713 | 1 | 1 |
| Cys/Ala | CTTTTAAACAAGAAATTTGCAATTTCATTAAAAAGTCCA | 9097..9132 (+) | 102 | 28 | - | - |
| Cys-2 | TACTTTTTAATGAAATTGCAAATTTCTTGTTTAAAAGTTACCA | 9094..9133 (-) | 1221 | 262 | - | - |
| Asp | TAACCATTTTTAGTTTTGTCAATACTTCTATAAGGTTAACCA | 2166..2204 (+) | 4047 | 583 | 0 | 0 |
| Glu | GAGTCTTTTTTTAGTGTTTTCAGCACTATTTTTTGACTTACCA | 4252..4291 (+) | 325 | 49 | - | - |
| Phe | AGTCAAAAAAATAGTGCTGAAAACACTAAAAAAAGACTACCA | 4253..4289 (-) | 4028 | 335 | - | - |
| Gly | TTCCTTACATTCTTATTTCCAATTAGATTAATAGGGAAACCA | 3786..3821 (+) | 2816 | 502 | 0 | 0 |
| His | AGATTTTTAAAACAGTCTGTGGAACTGAAGAAAAGTCTACCA | 5946..5984 (-) | 4459 | 356 | 3 | 1 |
| Ile | TGTTCTTTTTAACACTCTGATAAGGTGTAAATTTGAGCAACCA | 10067..10104 (+) | 806 | 245 | 11 | 1 |
| Lys | AGGGCATATATTTAGTAGGTGTCTTTTAAATACCTTGACCAACCCTACCA | 11249..11295 (+) | 9780 | 1255 | 0 | 0 |
| Leu1 | AATACATATTTTTGATTTAGGATTGAAATTTGAGTATTACCA | 12886..12923 (-) | 555 | 59 | 0 | 0 |
| Leu2 | CTTAACATTTTTTTAGAATTTAAGATTCTAATTTGCTTGAGCTTGTGTTAAGACCA | 4288..4340 (+) | 3085 | 1792 | 5 | 0 |
| Met | CCAGACTATTTAGGATTCATAACCCTTATTAAAACTGGACCA | 10106..10144 (+) | 3915 | 56 | 92 | 17 |
| Asn | TTTCATTAATCTTGCTTTGTTAGAGCTTAAAATTTGTGGAAACCA | 12822..12860 (-) | 1405 | 222 | 5 | 1 |
| Gln | TGGGATATTTTAAAATTCACTTTTTGGAAGTGAAAGTGTTTCCAACCA | 10141..10184 (-) | 1718 | 106 | 31 | 3 |
| Arg | TTGGTAAATAAGTAGTTTCGGCCTTCTAAACCAAACCAAACCA | 12792..12824 (+) | 2876 | 283 | 301 | 64 |
| Ser1 | AGAGTGATTAAAGAACTTCTAATTCTTGAAAGATTCTATCACACTCTTCCA | 4200..4247 (+) | 9010 | 2548 | 38 | 14 |
| Ser2-1 | GTTATTTTATAGAACTTTGAAAGTTCTCTTTTAGTAACTCCA | 9060..9098 (+) | 619 | 301 | 1 | 0 |
| Ser2-2 | AAACTTTTATTTAGTGTTTTGAAGACACTAGGAAGACCTGAAGTTTACCA | 10254..10299 (-) | 931 | 233 | - | - |
| Thr | TTTCTAACATTTGGATTTTGTAAATCCTAAATAAAGAAAACCA | 7522..7560 (+) | 1846 | 39 | 11 | 20 |
| Val | TTAACCTTATAGAAGTATTGACAAAACTAAAAATGGTTAAACCA | 2165..2204 (-) | 27 | 1 | 0 | 0 |
| Trp | GAAACTTCAGGTCTTCCTAGTGTCTTCAAAACACTAAATAAAAGTTTCACCA | 10253..10301 (+) | 3341 | 37 | - | - |
| Tyr | GGTGTTTTAAGTAAATTGTAAATTTAATTTAAAAACCACCA | 10185..10214 (-) | 472 | 139 | 0 | 0 |
| Pro | CGAGAGTAATTCTGCATTGGGGGTAGTTTTTTGTCTCGACCA | 10215..10253 (-) | 1662 | 337 | 30 | 10 |
